# Substrate recognition by human separase

**DOI:** 10.1101/2025.03.24.644901

**Authors:** Jun Yu, Sophia Schmidt, Margherita Botto, Kitaik Lee, Chloe M. Ghent, Jonah M. Goodfried, Andrew Howe, Francis J. O’Reilly, David O. Morgan, Andreas Boland

## Abstract

The protein complex cohesin encircles the sister chromatids in early mitosis^1^. At anaphase onset, sister separation is triggered by cleavage of the cohesin subunit SCC1/RAD21 by the cysteine protease separase^2–5^. SCC1 contains two cleavage sites, where cleavage is stimulated by SCC1 phosphorylation^5,6^. The molecular mechanisms of substrate recognition and cleavage are only partly understood^7^. Here, we determined a series of cryoEM structures of human separase in apo-or substrate-bound forms that, together with biochemical analysis, provide novel insights into the regulation of separase cleavage activity. We verify the first SCC1 cleavage site and reassign the second site. We show that multiple substrates, including separase autocleavage sites^8,9^ and the two SCC1 cleavage sites, interact with several docking sites in separase, including four phosphate-binding sites. We also describe the structural basis of the interaction between the cohesin subunit SA1/A2 and separase, which promotes cleavage at the second site in SCC1. Finally, using cross-linking mass spectrometry and cryoEM, we propose a model of how cohesin is targeted by human separase. Our work provides an extensive functional and structural framework that explains one of the most fundamental events in cell division.

## Introduction

In early mitosis, the sister chromatids are linked by the protein complex cohesin^1,10^. Chromosome segregation in anaphase is initiated by the proteolytic cleavage of the SCC1/RAD21 subunit of the cohesin ring by the cysteine protease separase^2–5^. Human separase is a large protein (2120 aa) composed of three domains: an N-terminal HEAT-repeat domain, a central TPR-like domain, and a C-terminal caspase-related protease domain^11,12^. The central TPR-like domain contains two large unstructured inserts and several smaller loop regions, including three separase autocleavage sites^8,9^ in insert 2 (**Fig. 1a**, top).

**Figure 1.**
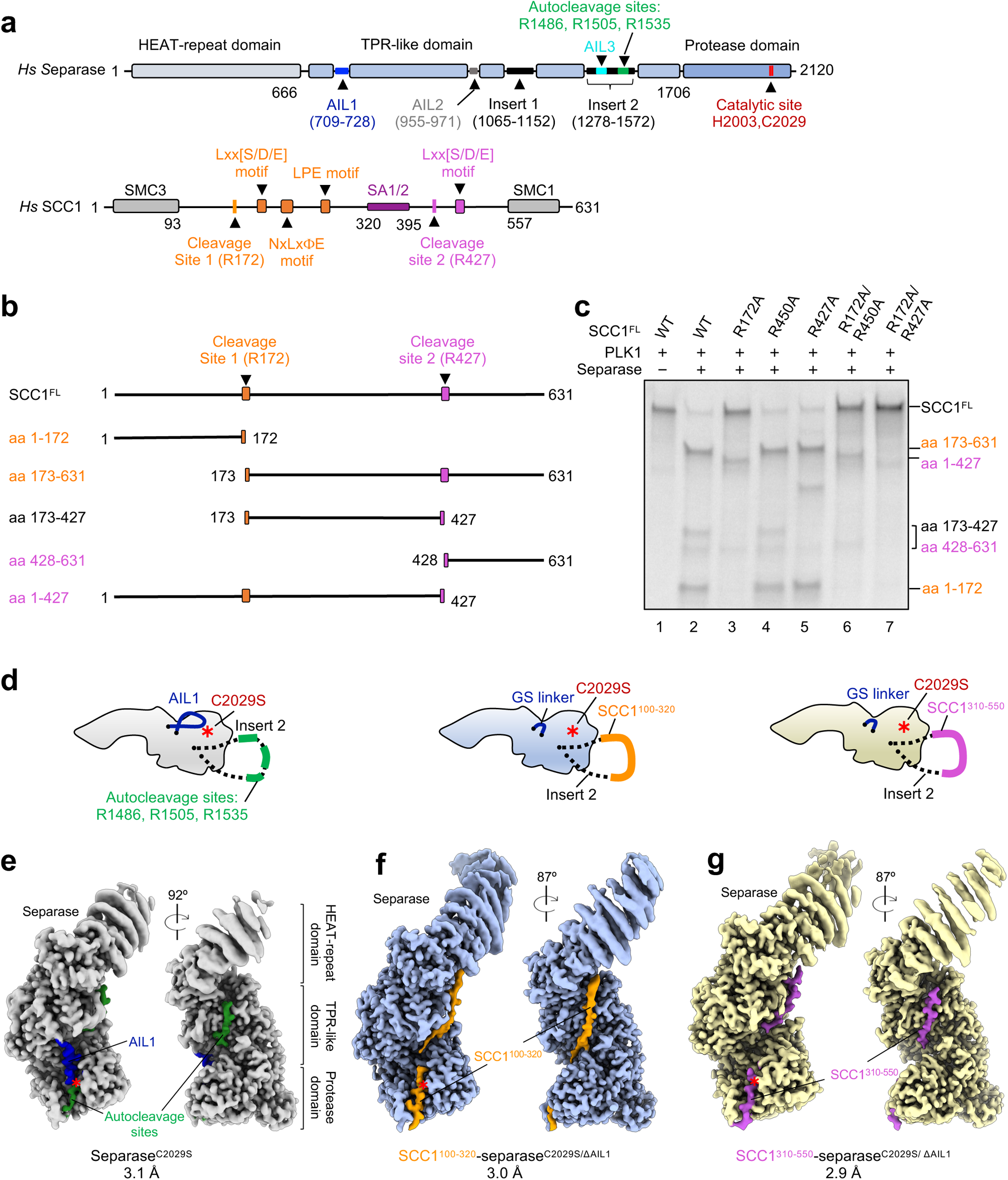
Cleavage of SCC1 at two sites and cryoEM reconstructions of human separase in apo or substrate-bound states. **a**, Domain organization of human separase and SCC1. Separase consists of three domains shown as blocks: a HEAT-repeat domain (grey), a TPR-like domain (light blue) and a C-terminal protease domain (blue). The TPR-like domain contains auto-inhibitory loops 1-3 (AIL1-3), shown with lines in dark blue, grey and cyan, as well as two large flexible insertions, insert 1 (aa 1065-1152) and insert 2 (aa 1278-1572) illustrated in black. Three autocleavage sites (green line) are in the insert 2. The catalytic site consists of two conserved residues His2003 and Cys2029 (red). The N-terminal SMC3-binding domain and C-terminal SMC1-binding domain of SCC1 are indicated with grey blocks. SCC1 contains two cleavage sites: cleavage site 1 (R172) with nearby substrate motifs (orange) and cleavage site 2 (R427) with nearby substrate motifs (orchid). The SA1/2-binding region of SCC1 is shown in purple. **b,** Schematic representation of SCC1 cleavage products resulting from two sites. **c,** Cleavage assay using ^35^S-labelled wild-type SCC1 and mutants as substrates. **d,** Fusion strategy for reconstitution of separase-substrate complexes. Left, inactive apo-separase (Separase^C2029S^, light grey) with insert 2 shown with a dashed line. AIL1 and autocleavage sites are color-coded as in **a**. Middle and right, region containing three autocleavage sites in insert 2 is replaced by SCC1 fragments: SCC1 site 1 (aa 100-320, orange) or SCC1 site 2 (aa 310-550, orchid). AIL1 is replaced by a GS linker (ΔAIL1)^11^. The C2029S mutation in separase indicates an inactive variant. Red star, the catalytic site. **e-g,** CryoEM reconstructions of apo separase (C2029S, **e**) and separase bound to SCC1 (aa 100-320, **f**) and SCC1 (aa 310-550, **g**). The color codes are the same as in **d.**

Due to its central function in cell cycle progression, separase activity is tightly controlled by multiple mechanisms. In vertebrates, activity is regulated through the concerted action of two mutually exclusive inhibitors: the universal inhibitor securin^8,13,14^ and a heterotrimeric complex of cyclin-dependent kinase 1 (CDK1), its regulatory subunit CKS1 or CKS2, and the cyclin subunit B1 or B2 (the CCC complex)^15–17^. Recent cryoEM structures provided detailed insights into the mechanisms of human separase inhibition^11^. Securin directly blocks substrate access to the active site and nearby docking sites by mimicking substrate binding, whereas the CCC complex rigidifies autoinhibitory loop segments (AILs) in separase that sterically preclude substrate binding.

Cleavage of separase substrates occurs at [D/E/S]ΦExxR motifs (P6–P1 substrate positions, where cleavage occurs after the invariant R at the P1 position)^2,5,18^. The structural basis of cleavage site binding at the active site was first revealed by the crystal structure of the separase protease domain from *Chaetomium thermophilum* fused to a short peptide mimicking the minimal SCC1 cleavage motif^7^. Two cleavage sites have been described in human SCC1^5,6^. R172^SCC1^ is well established as cleavage site 1^5,11,19^. Site 2 is not as well understood, but it has been proposed^6^ that cleavage occurs after R450^SCC1^.

Multiple ExxR sequences exist in SCC1 and across the proteome that are not cleaved, indicating the need for additional specificity determinants. Recently, docking sites outside the active site (exosites) were identified by biochemical and structural studies^11,19,20^. SCC1 contains LPE and NHLEYE motifs that are required for efficient site 1 cleavage, and securin contains related motifs required for binding. The cryoEM structure of the separase-CCC complex revealed that autoinhibitory loops dispersed throughout separase block substrate-binding sites. These lines of evidence suggest that substrates interact at a series of docking sites along the surface of separase, from the C-terminal active site to the N-terminal HEAT-repeat domain.

Cleavage of substrates by separase is often enhanced by substrate phosphorylation. Best understood is phosphorylation at the P6 position^7,21,22^. The phosphoserine in the pSxExxR motif is ideally positioned to interact with a large basic patch next to the active site^7,12^. But there are other, more distant phosphorylation sites that stimulate cleavage. In SCC1, phosphorylation of multiple sites enhances cleavage^6^. Cleavage of other substrates (Rec8 and PCNT) is also stimulated by phosphorylation at multiple sites^23,24^. However, we do not understand the structural basis for separase interaction with these phosphorylation sites.

Importantly, we also do not know how separase interacts with the complete cohesin complex. Cohesin is a ring-shaped protein complex composed of four protein subunits^25^: the Structural Maintenance of Chromosomes (SMC) proteins SMC1 and SMC3, the kleisin subunit SCC1, and the SCC3 subunit (SA1 or SA2 in mammals), which binds the disordered region of SCC1 between the two separase cleavage sites^26,27^. It is conceivable that subunits other than SCC1 interact with separase to influence SCC1 cleavage.

We employed a structural approach to deepen our understanding of substrate recognition and cleavage by human separase. Using cryoEM structures of separase in apo– and substrate-bound forms, we define the human SCC1 cleavage sites and provide insights into the recognition of substrate motifs and phosphorylation sites distant from the cleavage site. We also describe an interaction between the cohesin subunit SA2 and separase, which promotes SCC1 cleavage at site 2. Finally, we propose a model of separase binding to the cohesin ring.

## Results

### Identification of separase cleavage sites in human SCC1

We first set out to corroborate the two described^5,6^ cleavage sites in human SCC1 *in vitro*. We purified active separase and analysed separase-mediated cleavage of radiolabelled wild-type and mutant forms of SCC1 in established cleavage assays^11,19^. Because phosphorylation of SCC1 stimulates its cleavage by separase^5,6,11,21,22^, the purified kinase domain of PLK1^28^ was routinely added to cleavage reactions.

We found that full-length phosphorylated SCC1 is efficiently cleaved by separase, resulting in a pattern that is consistent with cleavage at two sites (**Fig. 1b** and **Fig. 1c**, lane 1 vs lane 2). Mutation of the previously established site 1, R172, resulted in a substantial decrease in SCC1 cleavage (**Fig. 1c**, lane 3). In contrast, mutation of R450, the second proposed cleavage site, did not affect cleavage (**Fig. 1c**, lane 2 vs lane 4)^5,6^. To identify the second SCC1 cleavage site, we generated a series of ExxR mutants in SCC1 and found that mutation of R427 strongly impaired the generation of smaller cleavage products (aa 173-427 and aa 428-631) (**Fig. 1b** and **Fig. 1c**, lane 5), even though cleavage of full-length SCC1 appeared unaffected. A double mutant containing mutations of both R172 and R450 showed a phenotype identical to that of the R172-only mutant, whereas a R172 R427 double mutant was not cleaved by separase (**Fig. 1c**, lane 6 and 7). These results suggest that the first cleavage site in human SCC1 is R172 as reported, but the second site is at R427 and not R450.

### Structure of inactive apo-separase reveals binding sites for autocleavage sites

The structure of separase in the absence of bound inhibitor proteins has not yet been determined for any species^7,11,29,30^. To gain further insights into substrate cleavage by human separase, we subjected purified wild-type and inactive mutant (catalytic cysteine replaced with serine: separase^C2029S^) apo-separase to cryoEM analysis (**Fig. 1 d, e** and **Extended Data Figs. 1** and **2**). Inactive separase is characterized by a single band on SDS-PAGE, whereas active separase autocleavage results in multiple distinct cleavage products (**Extended Data Fig. 1a, b**). We determined the structures of active and inactive apo-separase at overall resolutions of 3.3 and 3.1 Å, respectively (**Extended Data Fig. 1i-l, Extended Data Fig. 2** and **Extended Data Table 1**). Separase consists of an N-terminal HEAT-repeat domain, a central TPR-like domain and a C-terminal protease domain (**Fig. 1a**, **e**). As observed in previous human separase structures^11^, the N-terminal HEAT-repeat domain of separase moves as a flexible rigid body with respect to the rest of the protein (**Fig. 1e** and **Extended Data Fig. 1i, j**). Overall, the structures of cleaved active and intact inactive apo-separase are indistinguishable, with an RMSD (root mean square deviation) of 0.524 Å over 1089 C_α_-atoms, confirming previous evidence that the autocleavage products of active separase remain tightly associated^8^. The apo-separase structures are also almost identical to published human separase structures^11^ bound to the inhibitors securin (PDB: 7NJ1) or the CDK1-cyclin B 1-CKS1 (CCC) complex (PDB: 7NJ0), or when bound to the substrate SCC1 (this study, **Extended Data Fig. 3** and **Extended Table 2**). Thus, the overall separase structure is not affected by interactions with inhibitors or substrates.

In the EM maps of inactive and active apo-separase, we observe clear densities for the autoinhibitory loop segment AIL1, which we identified in our previous studies of the separase-CCC complex^11^. AIL1 is positioned in a substrate-binding cleft next to the catalytic site (**Fig. 1d, e** and **Extended Data Figs. 4a, b**). Deletion of AIL1 increases separase activity through increased substrate affinity^11^. Furthermore, the high-affinity pseudosubstrate securin contains a motif that displaces AIL1. In the apo-separase structures, however, the densities for the AIL1 segments are weaker than those of the separase-CCC structure, probably due to a lower occupancy of AIL1 in this conformation in the apo-separase EM maps. It therefore seems likely that the AIL1 segment exhibits inherent flexibility and acts as a modulator of separase cleavage activity. When bound to native substrates or securin the loop is displaced, which allows positioning of the binding partner in the active site pocket.

In addition to the density of AIL1, the EM map of inactive apo-separase (separase^C2029S^) contains other densities around the active site and at a conserved positively-charged patch in the central TPR domain (**Fig. 1e** and **Extended Data Fig. 4b**). This density is absent in the EM maps of active apo-separase or inactive separase with the three autocleavage sites deleted (**Extended Data Figs. 4a** and **5**), suggesting that these densities are related to autocleavage. The quality of the separase^C2029S^ map allows the unambiguous sequence assignment of these densities. The density around the active site represents the first autocleavage motif ^1481^GP**E**IM**R**^1486^, whereas the second additional density can be annotated to the third autocleavage motif ^1530^EW**E**LL**R**^1535^ (P6-P1 position with the ExxR cleavage motif in bold)^8^. This second binding site on separase was previously predicted using AlphaFold2^32^ structure predictions of yeast separase in complex with SCC1 or Rec8^20^. It is likely that the first autocleavage site inserted into the catalytic pocket represents the primary cleavage site, but cleavage can also occur at the two other autocleavage sites^8,9^. As we noted recently, the first autocleavage site of separase overlaps with a potential PP2A binding site, and autocleavage of separase abrogates PP2A binding^12,33^. The separase-binding motifs are discussed in detail in a later section.

### Overall architecture of separase-SCC1 complexes

We next sought to determine the structure of SCC1 bound to separase. Attempts to solve a separase-SCC1 structure using wild-type but inactive proteins were unsuccessful, even when full-length SCC1 was fused to the N-terminus of separase and autocleavage sites in insert 2 were removed (**Extended Data Fig. 5**). We therefore designed new SCC1-separase^C2029S^ fusion constructs based on insights obtained from the structures of apo-separase, separase bound to the CCC complex, and the SCC1-separase fusion protein. To reduce the substrate-blocking effect of AIL1, we replaced this loop with a short flexible linker as previously described^11^. We replaced the autocleavage sites of insert 2 with defined SCC1 fragments containing either cleavage site 1 (SCC1^100–320^) or cleavage site 2 (SCC1^310–550^) (**Fig. 1d, middle and right**). We then purified these new SCC1-separase fusion constructs (**Extended Data Fig. 6a, b)** and incubated them with PLK1 prior to cryoEM analysis. We determined the structures of SCC1^100–320^ (site 1) or SCC1^310–550^ (site 2) bound to separase at overall resolutions of 3.0 and 2.9 Å, respectively (**Extended Data Fig. 6c-j, Extended Data Fig. 7**, and **Extended Data Table 1**). The two structures show that each fragment binds similarly to separase, using the same binding sites to zigzag across the surface of the enzyme in an antiparallel and extended form (**Fig. 1f, g** and **Fig. 2a**), reminiscent of the inhibitor securin^11,29,30^.

**Figure 2.**
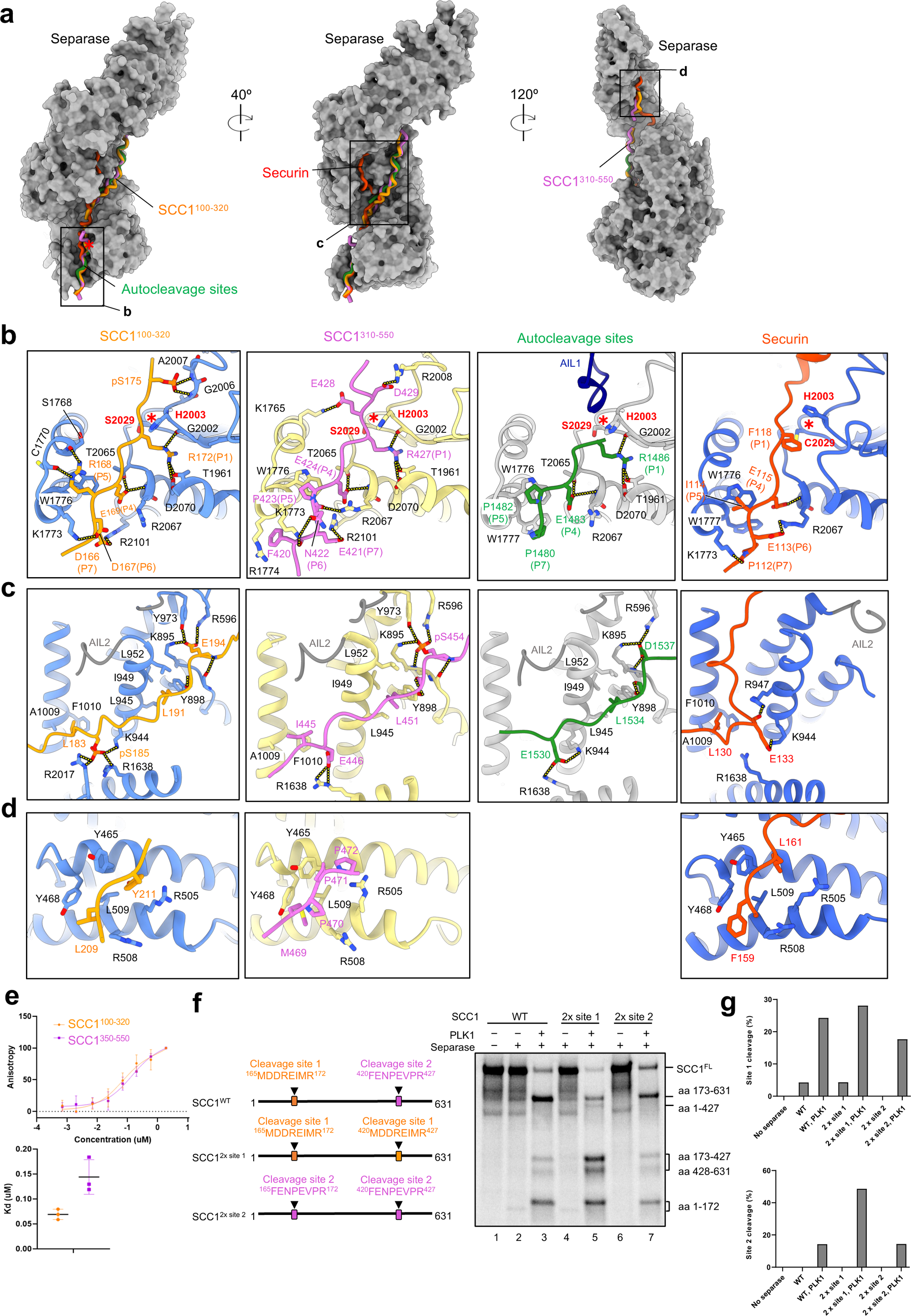
Multiple substrate motifs interact with separase. **a**, Views of SCC1 site 1 (aa 100-320) and site 2 (aa 310-550), plus autocleavage sites and securin binding across the three domains of separase. Structures of Separase^C2029S^, securin-separase complex (PDB: 7NJ1) and SCC1^310–550^– separase^C2029S/ΔAIL1^ are aligned to SCC1^100–320^–separase^C2029S/ΔAIL1^. Separase is depicted as surface representation in dark grey. SCC1 (orange or orchid), securin (orange red) and autocleavage sites (green) are shown as cartoon. **b,** Close-up view of SCC1, securin and autocleavage site binding near the catalytic site of separase. Hydrogen bonds are indicated with yellow dashed lines. **c,** Close-up view of SCC1, securin and autocleavage site binding to the TPR-like domain of separase. **d,** Close-up view of SCC1 and securin binding to the HEAT-repeat domain of separase. **e,** Affinity measurement of SCC1^100–320^ and SCC1^350–550^ binding to Separase^C2029S^ using fluorescence polarization. Each experiment was repeated three times; data points indicate mean +/-SEM. **f,** Cleavage assay of ^35^S-labelled wild-type SCC1 and mutants containing two copies of cleavage site 1 motif (SCC1^2x^ ^site^ ^1^) or two copies of cleavage site 2 motif (SCC1^2x^ ^site^ ^2^). Left, schematic representation of wild-type SCC1 and mutants. Right, autoradiograph of SCC1 cleavage. Results are representative of three independent experiments. **g,** Top, quantification of relative abundance of site 1 cleavage fragment (aa 1–172). Below, quantification of relative abundance of site 2 cleavage fragments (aa 173–427 plus aa 428–631). For normalization, the intensities of the cleavage fragments were divided by the total intensities of all bands in the respective lanes.

While binding of the SCC1 fragments overlaps with securin around the catalytic site and the N-terminal HEAT-repeat domain of separase, the binding paths differ at the TPR-like domain (**Fig. 2a-d**). Binding of securin to the TPR-like domain prompts a movement of the autoinhibitory loop 2 (AIL2) from its regular position (as observed in the apo-separase structure) to a binding site that is usually recognized by substrates. Securin therefore binds to different TPR-like repeats than substrates. Conversely, when in the apo or substrate-bound state, AIL2 of separase adopts a conformation incompatible with securin binding. Therefore, AIL2 reinforces mutually exclusive binding of substrates or inhibitors where the binding paths do not overlap. The different binding modes likely result in different separase binding affinities (**Extended Data Fig. 3b, c**).

### Four positively-charged docking sites anchor substrate binding to separase

The structure of cleavage site 1 bound to separase shows that R172 of SCC1 (P1 position) is deeply inserted into the catalytic pocket of separase, thereby orchestrating a configuration of the active site that allows substrate cleavage (**Fig. 2b**, SCC1^100–320^ and **Extended Data Fig. 9c** below). A hydrogen-bonding network between R172^SCC1^ and residues lining the catalytic pocket, such as D2070, T1961 or G2002, facilitates a rotamer torsion of H2003^Separase^ that, as a result, is optimally positioned for catalysis. This substrate-induced cleavage mechanism coordinated by an invariant arginine in substrates has been conserved in evolution^7,29,34^ (**Extended Data Fig. 8a**). Substrate binding at the P1-P6 positions near the active site is mediated by multiple interactions. R168^SCC1^ stacks onto W1776^Separase^ at the P5 position and hydrogen bonds through its guanidinium group with the main chain oxygens of S1768^Separase^ and C1770^Separase^.

Critical electrostatic interactions form between D166^SCC1^ and E169^SCC1^ (P7 and P4 positions, respectively) and a positively-charged patch immediately upstream of the separase active site. Key residues include R2067, R2071 and R2101 (**Fig. 2b**, SCC1^100–320^ and **Extended Data Fig. 9c**). This conserved positive patch explains how negatively charged residues usually conserved at the P7 and the P6 substrate positions are recognized, and how phosphorylation at the P6 position stimulates cleavage of SCC1 in budding yeast^7,18,21^ and likely the human kinetochore protein Meikin^35^. We refer to this patch as phosphate-binding site 1 (P-site 1).

Although human SCC1 is not phosphorylated at or near the P6 position (**Extended Data Fig. 8a**), phosphorylation elsewhere has been described to stimulate cleavage of site 1^6^. The structural basis of this effect is not known. In our structure of PLK1-phosphorylated SCC1^100–320^–separase^C2029S/ΔAIL1^, we found that serines 175 and 185 of SCC1 are (partially) phosphorylated, according to EM densities and mass spectrometry (**Fig. 2b, c** and **Extended Data Figs. 4c, f, g** and **9a-d**). Interestingly, phosphorylated S175^SCC1^ inserts in a pocket next to the active site, where it would sterically clash with the catalytic histidine H2003 in a securin-induced inactive rotamer conformation (**Extended Data Fig. 10a**). Thus, phosphorylation of S175^SCC1^ promotes the active rotamer conformation. The ideal positioning of H2003 is further supported by pS175^SCC1^ forming multiple hydrogen bonds with the side chain of R2008^Separase^ and the main chain amino groups of G2004^Separase^, A2005^Separase^ and G2006^Separase^, forming a second phosphate-binding pocket (P-site 2; **Fig. 2b** and **Extended Data Fig. 9b, c**). The latter group of residues are in a loop segment adjacent to the catalytic histidine that is likely to be rigidified upon binding to pS175^SCC1^. Indeed, as observed previously^6^, mutation of S175^SCC1^ to alanine (S175A) mildly reduces separase cleavage activity and a S175E mutant shows slightly enhanced cleavage (**Extended Data Fig. 10b**).

The mild effects of S175 mutations suggested that additional SCC1 phosphosites stimulate separase cleavage activity. Indeed, we observe a density for phosphorylated serine 185 in our EM map (**Extended Data Fig. 4c, g**). pS185^SCC1^ is recognised by a strongly positively-charged third phosphate-binding site, P-site 3, created by K944, R1638, and R2017 at an interdomain cleft of separase (**Extended Data Fig. 9d**). L183^SCC1^ is inserted into an adjacent hydrophobic groove and stabilizes pS185 at the phosphate-binding site (**Fig. 2c**). This site is also bound by a previously described LPEE motif in securin^19^, in which the leucine binds the hydrophobic groove and the second glutamate is recognised by the positively-charged pocket (**Fig. 2c**). We mutated the ^185^STTTS^189^ motif, which has been shown to be phosphorylated in mitosis^6^, and studied its effects in combination with the S175A^SCC1^ mutation. The combination of multiple mutations noticeably reduced cleavage efficiency by separase (**Extended Data Fig. 10c**).

The TPR-like domain contains a fourth positively-charged site, P-site 4, that in the separase-securin complex structure is blocked by AIL2 (**Extended Data Fig. 3b, c**). When SCC1 site 1 is bound, AIL2 is in an open conformation as in the apo-structure, enabling access to a positive patch recognized by E194^SCC1^ (**Fig. 2c** and **Extended Data Fig. 9d**). E194^SCC1^ hydrogen bonds with R596, K895 and Y973 of separase. A highly conserved leucine residue (L191^SCC1^), three amino acids N-terminal of E194^SCC1^, binds to a hydrophobic pocket and stacks onto Y898 of separase to serve as an additional anchor point. The resulting Lxx(S/D/E) motif is highly conserved in separase substrates (**Extended Data Fig. 8b**). Interestingly, it is also mimicked by AIL2 (L959 and D972) (**Extended Data Fig. 3c**). In our SCC1 site 2 structure, the acidic side chain is replaced by a phosphorylated serine as described below, and thus P-site 4 represents a fourth phosphate-binding site.

Lastly, we also observe binding of an SCC1 fragment to the N-terminal HEAT-repeat domain of separase. This interaction is mediated by hydrophobic contacts: L209^SCC1^ and Y211^SCC1^ mediate stacking interactions with Y465^Separase^ to tether SCC1 site 1 to the separase HEAT-repeat domain (**Fig. 2d** and **Extended Data Fig. 4e**). These residues are part of an NHLEYE motif (**Extended Data Fig. 8c**) that we described previously as an enhancer of SCC1 cleavage^11^. Due to lower resolution in this map region, model building was aided by AlphaFold3 predictions^31^.

Our previous work showed that an LPE motif in SCC1 is required for site 1 cleavage in the absence of phosphorylation^19^. As mentioned above, a similar LPEE motif in securin interacts with P-site 3 of separase^19^. However, in the separase-SCC1 structure we observe that the LPE motif is not involved in binding to P-site 3 when SCC1 is phosphorylated. Indeed, we found that cleavage of phosphorylated site 1 in vitro does not require the LPE motif (**Extended Data Fig. 9e**).

Using the same fusion strategy, we determined the structure of separase bound to SCC1 cleavage site 2 (SCC1^310–550^–separase^C2029S/ΔAIL1^). Overall, the binding mode of the second cleavage site is analogous to that of site 1. R427 of SCC1 is inserted into the catalytic site pocket, consistent with our biochemical identification of R427^SCC1^ as the second cleavage site (**Fig. 2b**, SCC1^310–550^ and **Extended Data Fig. 9c**). E421^SCC1^ and E424^SCC1^ (P7 and P4 positions, respectively) are recognised by residues forming P-site 1 (R2067, R2071 and R2101), whereas W1776^Separase^ forms stacking interactions with P423^SCC1^ at the P5 position. In addition, F420^SCC1^ stacks onto R1774^Separase^. The phosphoserine 175 described for site 1 is substituted with D429, which possibly hydrogen bonds with R2008^Separase^, part of the loop that harbours the catalytic H2003^Separase^ (**Fig. 2b**, SCC1^310–550^). Similarly, phosphoserine 185 in site 1 is replaced with a glutamate, E446^SCC1^, that binds to the basic residues of P-site 3 (K944, R947, R1638 and K1645). Another similarity between site 1 and site 2 is that E446^SCC1^ binding to P-site 3 is supported by a hydrophobic N-terminal anchor: I445^SCC1^ binds to the adjacent hydrophobic groove, thereby stabilizing and precisely positioning E446^SCC1^. While P-sites 2 and 3 recognize phosphoserines in cleavage site 1 but not in site 2, the situation is reversed for P-site 4. Here, as described above, site 1 binds to separase through E194, whereas at site 2 we observe a phosphoserine bound to P-site 4 (**Fig. 2c**, SCC1^310–550^ and **Extended Data Fig. 9b, d**). The phosphate group of pS454^SCC1^ forms multiple hydrogen bonds with R596, K895 and Y973 of separase. Individual or combined mutations of I444^SCC1^, E446^SCC1^, L451^SCC1^ or S454^SCC1^ to alanines impairs the generation of site 2 cleavage products in vitro, indicative of reduced binding of SCC1 to separase (**Extended Data Fig. 10c**). These results are consistent with previous evidence that a S454A mutation inhibits cleavage at the second site^6^.

Comparable to site 1, SCC1 also binds via site 2 to the N-terminal HEAT-repeat domain. M468 and P470 of a conserved MPPP-motif (aa 468-471) tether SCC1 to the HEAT-repeat domain (**Extended Data Fig. 8c**). Here, M468^SCC1^ binds separase as a hydrophobic anchor like L209^SCC1^ of site 1. P470^SCC1^ of site 2 forms stacking interaction with Y468^Separase^ analogous to Y211^SCC1^ of site 1 (**Fig. 2d**, SCC1^100–320^ and SCC1^310–550^ and **Extended Data Fig. 4e**). Interestingly, this binding mode is mimicked by securin. F159^Securin^ and L161^Securin^ bind to the same HEAT repeat by hydrophobic and stacking interactions, respectively (**Fig. 2d**, Securin, and **Extended Data Fig. 8d**)^19^.

With SCC1 structures in mind, we can now understand in more detail the binding of the first autocleavage site in our structure of inactive apo-separase. In this structure, R1486^Separase^ is inserted into the active site, priming the catalytic dyad in a conformation suitable for substrate cleavage. For the first autocleavage site, only E1483^Separase^ at the P4 position is recognised by P-site 1. At the P5 and P7 positions, two proline residues, P1482^Separase^ and P1480^Separase^, form stacking interactions with the highly conserved W1776^Separase^ and W1777^Separase^, respectively (**Fig. 2b**, Autocleavage sites). The prolines are interspersed by G1481^Separase^. Therefore, in this binding mode, the usually negatively-charged P6 substrate position that is recognised by P-site 1 is replaced by G1481^Separase^, which provides a flexible hinge to enable stacking interactions of P1482^Separase^ with W1777^Separase^. This different binding mode might explain the sequence variability of the substrate binding motif at the P6 position (**Extended Data Fig. 8a**). Another feature that discriminates binding of the autocleavage site from substrates or securin binding is that AIL1 is not displaced by the autocleavage motif. As a result, AIL1 is present in our EM map and the derived model (**Figs. 1e** and **2b**, Autocleavage sites). The molecular interactions between E1530^Separase^ and P-site 3 are reminiscent of E446^SCC1^ of SCC1 site 2. Further downstream, an LxxD motif binds to P-site 4 as observed for the LxxE motif of site 1 or the LxxpS motif of site 2 (**Fig. 2c**).

Comparing the binding modes of SCC1 cleavage sites 1 and 2 reveals commonalities but also differences in their recognition by separase (**Extended Movie 1**), which might result in different binding affinities. In addition, we observe a strong preferential cleavage of site 1 in our cleavage assays (**Extended Data Fig. 10d**). To estimate binding affinities for the two sites separately, we purified two SCC1 fragments containing either cleavage site 1 or 2 (SCC1^100–320^ and SCC1^350–550^). Both fragments were N-terminally linked to the Maltose-binding protein (MBP) and a tetra-cysteine peptide was fused to the C-terminus for binding by a specific fluorophore (FlAsH-EDT_2_). This setup allowed us to perform fluorescent polarization experiments. Binding of inactive separase to SCC1 causes a slowed rotation (higher anisotropy) of the fluorescently labelled SCC1 due to a substantial mass increase. Our assay shows that the SCC1^100–320^ construct exhibits a roughly 3-fold lower K_d_ than the SCC1^350–550^ construct (**Fig. 2e**). To reveal the effect of the cleavage motif alone (P8-P1 positions) on cleavage activity, independent of other binding sites, we designed two artificial SCC1 constructs. In a SCC1^2xsite1^ construct, cleavage site 2 was substituted by site 1 to generate a construct containing two copies of site 1. Conversely, we replaced cleavage site 1 with site 2 to generate a construct containing two copies of site 2 (SCC1^2xsite2^) (**Fig. 2f**). As we observed previously, phosphorylation of wild-type SCC1 strongly stimulates cleavage (**Fig. 2f**). SCC1^2xsite1^ is more efficiently cleaved than wild-type SCC1, resulting in enhanced generation of smaller cleavage products (**Fig. 2f**, compare lane 3 vs 5). In contrast, SCC1^2xsite2^ is less efficiently cleaved than wild-type SCC1 or SCC1^2xsite1^ (**Fig. 2f**). The results are quantified in **Fig. 2g**. As expected, if enhanced binding affinity is caused by increased electrostatic interactions upon substrate phosphorylation, we observed that cleavage efficiency declines at increasing salt concentrations (**Extended Data Fig. 9f)**.

### The cohesin subunit SA1 or SA2 stimulates SCC1 cleavage at the second cleavage site

The long disordered central region of SCC1 includes a binding site for the cohesin SA1/2 subunit (**Fig. 1a**, bottom)^36^. Given the proximity of the SA1/2 binding site to site 2, we speculated that SA1/2 might affect separase cleavage activity. Consistent with this possibility, we found that pre-incubation of SCC1 with SA1 or SA2 stimulates the generation of smaller cleavage products, while not affecting larger cleavage products (**Fig. 3a** and **Extended Data Fig. 11a**). SA2 stimulates cleavage of SCC1 site 2 but not site 1 even in the absence of PLK1 (**Extended Data Fig. 11a**).

**Figure 3.**
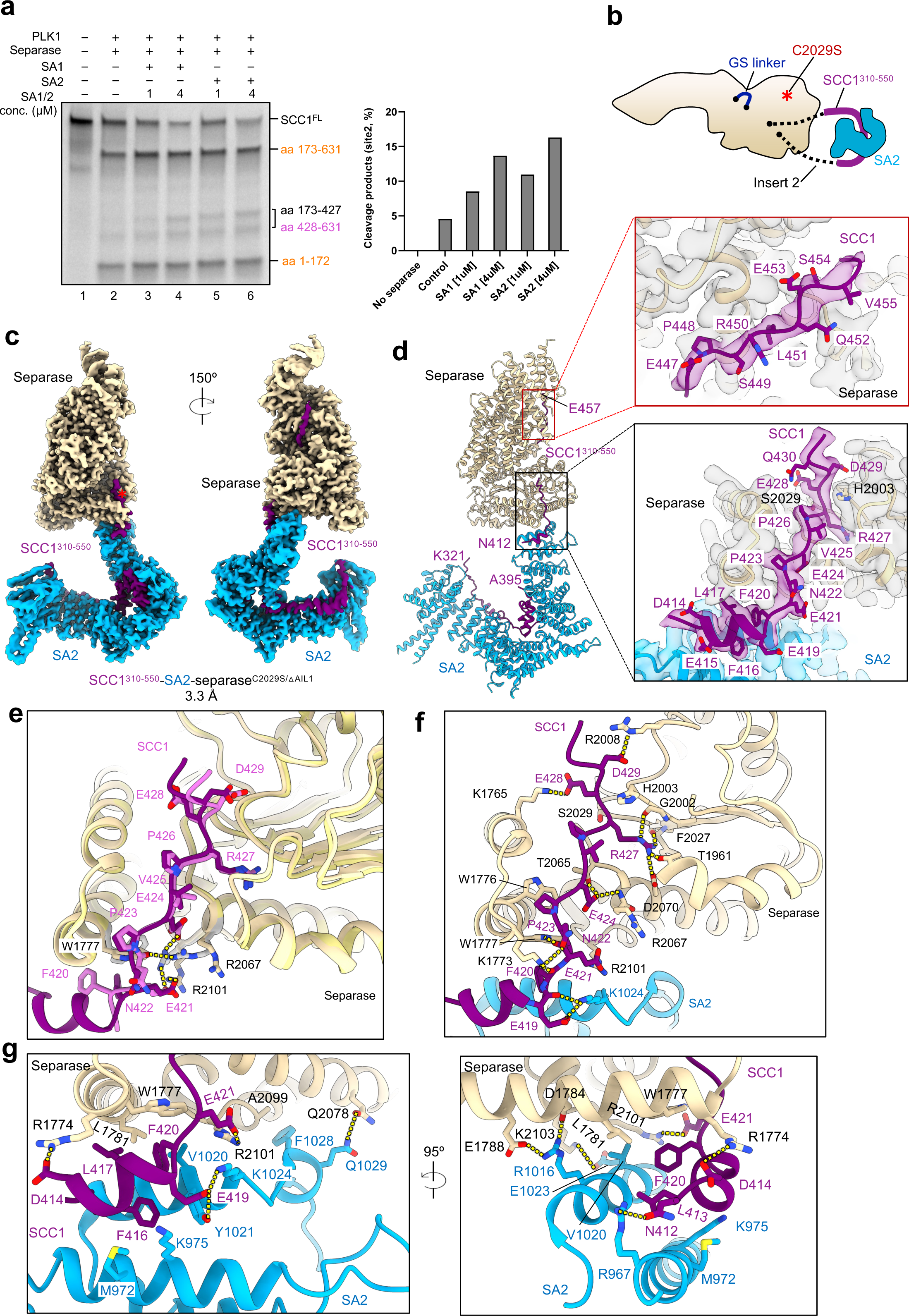
SA1/2 stimulates SCC1 cleavage at site 2 and cryoEM structure of separase bound to SCC1-SA2 complex. **a**, Cleavage assay showing stimulation of SCC1 cleavage at site 2 by SA1/2 proteins, tested at two different concentrations (1 and 4 µM). Left, autoradiograph showing cleavage products of ^35^S-labelled SCC1. Right, quantification of relative abundance of site 2 cleavage fragments (aa 173–427 and aa 428–631). For normalization, the intensities of the site 2 cleavage fragments were divided by the total intensities of all bands in the respective lanes. Results are representative of three independent experiments. **b,** Schematic representation showing the strategy for reconstituting SCC1-SA2-separase complex. SCC1 (aa 310-550) fragment (purple) includes both SA2 binding region and cleavage site 2. SA2 is shown in deep sky blue and separase is depicted in wheat. **c,** Views of the cryoEM map of the SCC1^310–550^–SA2-separase^C2029S/ΔAIL1^ complex. **d**, Ribbon representation of the SCC1^310–550^–SA2-separase^C2029S/ΔAIL1^ complex highlighting the interaction interfaces between the three proteins. Red box, cryoEM density showing the interaction interface between SCC1 and the TPR-like domain of separase. Black box, cryoEM density showing the interaction interface between SCC1, SA2 and the protease domain of separase. **e**, A close-up view showing SCC1 binding to the protease domain of separase in the complexes of SCC1^310–550^–SA2-separase^C2029S/ΔAIL1^ and SCC1^310–550^– separase^C2029S/ΔAIL1^. The two structures were aligned using separase as a reference and are shown in ribbon representation. SCC1 in the two complexes, depicted as sticks, is shown in purple and orchid, respectively. Side chains of W1777, R2067 and R2101 in the two complexes are shown in wheat and grey, respectively. Hydrogen bonds are indicated with yellow dashed lines. **f**, Close-up view of the cleavage site 2 motif of SCC1 binding near the catalytic site of separase. **g**, Two views of the interaction interface formed by a short helix of SCC1 (aa 413-419), C-terminal helices of SA2 and the protease domain of separase.

To explore these effects in detail, we prepared a complex composed of SCC1^310–550^– separase^C2029S/ΔAIL1^ and full-length SA2. The proteins formed a stable complex in vitro that was subjected to structural studies, without prior phosphorylation (**Extended Data Fig. 12a**). In our cryoEM analysis, however, we observed multiple classes: a ternary complex consisting of SCC1^310–550^– separase^C2029S/ΔAIL1^ and full-length SA2; SCC1^310–550^–separase^C2029S/ΔAIL1^; and SCC1^310–550^ bound to SA2, and determined their structures to resolutions of 3.4 Å, 2.8 Å and 2.9 Å, respectively (**Extended Data Fig. 12b-j** and **Extended Data Table 1**). Because of the heterogeneity of the sample, likely caused by low-affinity binding between SA2 and separase, and in addition because of preferential orientation of particles, we collected more than 50,000 micrographs to obtain high-resolution structures. The complex processing scheme is depicted in **Extended Data Fig. 13.**

The SCC1^310–550^–SA2-separase^C2029S/ΔAIL1^ complex is highly elongated and measures more than 250 Å in its longest and less than 50 Å in its shortest dimension (the flexible N-terminal HEAT-repeat domain of separase is not shown in **Fig. 3c, d**). The C-terminal HEAT repeat of the hook-shaped SA2 protein packs onto helices that surround the active site of separase to form a minimal protein-protein interface that is driven mainly by electrostatic interactions. For example, R1016^SA2^ forms hydrogen bonds with E1788^Separase^ and D1784^Separase^ and E1023^SA2^ interacts with K2103^Separase^ (**Fig. 3c, g**). This interface provides additional anchor points for SCC1 to be ideally positioned in and around the catalytic site (**Fig. 3d** and **Extended Movie 2**).

SCC1 binding to the TPR-like domain is virtually identical in the absence or presence of SA2, even though we did not phosphorylate SCC1 with PLK1 in this sample (**Fig. 3d** and **Extended Data Fig. 11b**). However, simultaneous binding of SCC1 and SA2 to the protease domain of separase causes a reorganisation of several side chains that are part of P-site 1 recognising the P6 substrate position. This includes R2067^Separase^, R2071^Separase^ and R2101^Separase^ as well as the highly conserved W1777^Separase^ (**Fig. 3e** and **Extended Data Fig. 11c**). W1777^Separase^ flips to a rotamer configuration that allows stacking interactions with F420^SCC1^. In the absence of SA2 binding, F420^SCC1^ is likely solvent-exposed and flexible. Similarly, R1774^Separase^ shifts by about 3 Å compared to the SCC1^310–550^–separase^C2029S/ΔAIL1^ structure. This new side chain position facilitates hydrogen bonding between R1774^Separase^ and D414^SCC1^, as well as the main chain oxygen of E415^SCC1^. The extensive hydrogen bonding network that ensures correct positioning of the substrate arginine in the catalytic side is illustrated in **Fig. 3f**, which includes a short amphipathic helix of SCC1^413–419^ (**Extended Data Fig. 11d**). Additional interactions that stabilise a short SCC1 α-helix at this interface of the separase protease domain and SA2 are between F416^SCC1^, M972^SA2^ and K975^SA2^, as well as hydrogen bonding between E419^SCC1^ and K1024^SA2^ (**Fig. 3g**). The overall structure of SA2 when bound to separase and SCC1 is the same as that in a binary complex composed of SCC1^310–550^ and SA2 (**Extended Data Fig. 11e**). In line with our biochemical studies that demonstrate that both SA proteins can stimulate SCC1 site 2 cleavage by separase, the residues forming the interface between SA2 and separase are conserved in SA1 (**Extended Data Fig. 11f, g**).

### The separase-cohesin complex structure

To gain further insights into separase substrate recognition, we reconstituted a separase-cohesin complex using a protein engineering strategy like the one described above. Full-length SCC1 was inserted in place of the autocleavage sites of inactive separase, and AIL1 was deleted to enhance substrate binding affinity. We co-expressed the cohesin subunits SA2 and the SMC proteins SMC1 and SMC3 together with the SCC-separase fusion construct and reconstituted the separase-cohesin complex in vitro (**Fig. 4a** and **Extended Data Fig. 14a**). A DNA fragment was added to the preparation to enhance complex formation. However, initial EM analysis did not yield promising separase-cohesin 2D classes and therefore we stabilised the complex using crosslinking reagents (**Extended Data Fig. 14b**).

**Figure 4.**
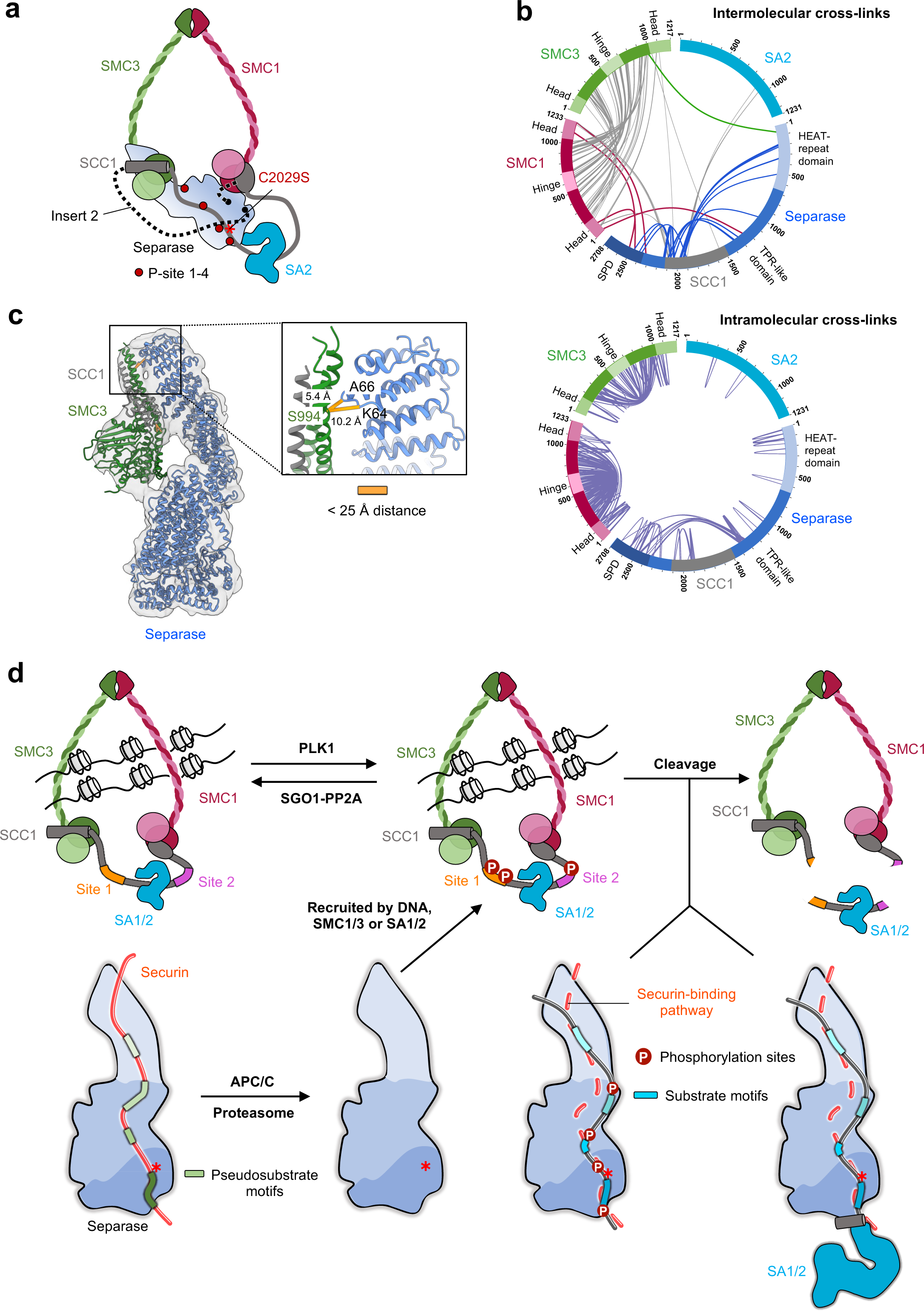
Cohesin targeting by separase is mediated by SMC subunits. **a**, Schematic representation of the strategy for reconstituting the cohesin-separase complex. Full-length SCC1 replaces the autocleavage sites in insert 2 (dashed line) of separase. SMC3 is colored in light and forest green, SMC1 in light and dark red, SCC1 in grey, SA2 in deep sky blue, and separase in a blue gradient **b,** Cross-linking mass spectrometry (XL-MS) analysis of cohesin-separase complex cross-linked using sulfo-SDA. Circular view showing intermolecular cross-links (top) and intramolecular cross-links (bottom) in the cohesin-separase complex. Each protein is color-coded as in **a**. Cross-links between the cohesin subunits (SMC1, SMC3 and SCC1) and separase are shown with dark red, green and blue lines, respectively. Intramolecular cross-links are shown in purple. **c,** CryoEM map and model of separase bound to SMC3-SCC1. Cross-linked residue pairs are indicated with orange lines. **d,** A model illustrating phosphorylation-or SA1/2-dependant regulation and the specificity determinants of cohesin cleavage by separase.

As a first step, we used the highly specific photoreactive crosslinker Sulfo-SDA to perform cross-linking mass spectrometry analysis (XL-MS). As expected, we observed numerous intra– and intermolecular crosslinks between the ATPase head domains and the long coiled-coil regions of SMC1 and SMC3 proteins (**Fig. 4b** and **Extended Data Fig. 14e**). Interestingly, we also detected several reproducible crosslinks between the SMC1 head domains and the C-terminal half of separase, as well as crosslinks between the SMC3 head domains and the N-terminus of separase (**Fig. 4b, c**).

We then used negative-staining electron microscopy to analyse a BS3-crosslinked sample. We detected three distinct complexes: the cohesin complex, separase bound to SA2, and separase bound to the head domains of SMC1 and/or SMC3 (**Extended Data Fig. 14c, d**). All separase-SMC1-SMC3 complex particles were then subjected to further 3D classification, which resulted in three different classes that showed additional densities beside the unambiguous separase density (**Extended Data Fig. 14f-h**). Class1 and class 2 showed additional density near the C-terminal half of separase, and the size of this extra density corresponded well to the SMC1 head domains. Combined with the evidence from XL-MS for crosslinks between SMC1 head domains and separase, we placed the SMC1 head domains in the C-terminal extra density (**Extended Data Fig. 14f, g**). Interestingly, the third class showed two extra densities: the C-terminal extra density plus an additional density of similar size near the separase N-terminus. Because we detected crosslink pairs between a helix protruding from the C-terminal head domain of SMC3 and the very N-terminus of separase, we placed the head domains of SMC3 in this second EM density (**Extended Data Fig. 14h)**.

We then subjected the cross-linked sample to cryoEM studies and resolved the complex structure at an overall resolution of 3.5 Å (**Extended Data Fig. 15)**. Again, we observed an extra density bound to the N-terminal HEAT-repeat domain (**Extended Data Figs. 14h** and **15d**). Using a mask on the N-terminal HEAT-repeat domain to select for particles with the SMC3 protein bound resulted in 26,747 particles that refined to an overall resolution of about 4.3 Å. However, the local resolution of the extra density next to the N-terminal domain is of lower resolution, indicating a high degree of flexibility of the complex (**Extended Data Fig. 15d**). We fitted the AF3-predicted model of the C-terminal SMC3 head domain in complex with SCC1 into this density, in agreement with our detected crosslinks (**Fig. 4c**).

Integrating all available data allows us to propose a model in which separase not only binds to the substrate SCC1 but also recognises the head domains of SMC1 and SMC3. In the case of SCC1 site 2 binding to separase, cleavage is further stimulated by an interaction with the SA1/2 subunit of cohesin, thereby providing additional SCC1 binding sites near the active site of separase (**Fig. 4a**).

## Discussion

Our studies reveal that separase substrates bind in an extended antiparallel orientation to a series of docking sites along the surface of separase, from its C-terminal active site to its N-terminal HEAT-repeat domain (**Fig. 4d**). These docking interactions enhance substrate affinity and specificity, focusing the enzyme on a very small number of specific cleavage sites among the many ExxR motifs that exist in SCC1 and the proteome. About 20% of proteins in the yeast proteome carry an ExxR motif, but none of these candidates other than SCC1 and Rec8 have been confirmed as a separase substrate^18^.

The specificity of substrate binding is likely explained by the presence of multiple positively-charged substrate-docking sites that can interact either with acidic side chains or with phosphorylated substrate residues. Interactions between phosphorylated side chains and these sites explain previous evidence that phosphorylation promotes cleavage of sites 1 and 2 in SCC1. We suspect that these phosphate-binding sites are also involved in the recognition of other substrates, such as Rec8 and PCNT, whose cleavage is promoted by multisite phosphorylation^23,24^.

Our studies of substrate binding, together with our previous analysis of the separase-securin complex, demonstrate that substrates and inhibitors employ a remarkably diverse array of modular sequence motifs to bind separase with high affinity and specificity. The ability of separase to interact with many different motifs allows interesting variations in the motifs employed by different binding partners and even by the same partner. For example, as mentioned above, the four positively-charged docking sites interact with phosphate in some cases and acidic residues in others, and P-site 1 can also accommodate other amino acids through a distinct binding mode. SCC1 site 1 can bind through two distinct mechanisms: unphosphorylated SCC1 depends on LPE and NHLEYE motifs for efficient cleavage of site 1 in vitro and in vivo^19^, but cleavage of phosphorylated SCC1 at site 1 does not require these motifs but instead depends on phosphorylated residues in SCC1. We further identify an important second motif, LxxpS/D/E, that is crucial for binding of both SCC1 cleavage sites as well as the separase autocleavage site. We conclude that the high affinity of substrate and securin binding depends on a diverse selection of multiple low-affinity sites, each of which can be modified slightly without greatly affecting overall affinity. Searches for combinations of these motifs across the proteome might lead to the identification of novel substrates.

It is critical for the success of mitosis that separase exhibits extreme specificity for a small group of substrates and avoids off-target cleavage of other proteins. One solution to this problem is that separase exhibits extremely low catalytic activity towards a short peptide substrate containing only the site 1 cleavage sequence^19^. Cleavage at a significant rate clearly requires the boost in affinity and activity that results from highly specific interactions between separase and its targets. These interactions depend on the binding of substrate sequence motifs, but it seems likely that additional mechanisms focus the enzyme on the correct sites. We show, for example, that specificity for site 2 is enhanced by an interaction between separase and the cohesin subunit SA1/2. Additional specificity for cohesin might arise from the ability of separase to interact with DNA^37^. Finally, our studies of a separase complex with complete cohesin complex suggest that additional interactions between separase and the SMC subunits are also be involved. The binding of the SMC head domains (from SMC1 and SMC3) might play an important role in cohesin recognition by interacting with separase on the opposite side from SCC1. These and other mechanisms ensure that the potentially dangerous protease activity of separase is focused precisely on the correct targets, resulting in a successful completion of cell division. Such mechanisms might also be exploited by future targeted drug design studies^38^. It is, for example, conceivable that downregulating separase activity in specific cancer types could be facilitated by a specific inhibitor cocktail that targets multiple substrate-docking sites on separase.

## Materials and Methods

### Expression vector construction

The cDNAs encoding human separase, SMC1A, SMC3 and STAG1/2 (SA1/2) were synthesized as gene-optimized versions for expression in *Spodoptera frugiperda* (Sf9) cells. The cDNA for SCC1 was PCR-amplified from a human cDNA library. The apo-separase plasmid was generated by subcloning synthesized cDNA (Thermo Fisher Scientific) of separase (aa 1-2120) into a pF1 vector with a Twin-StrepII tag at the C-terminus and 3xFLAG tag at the N-terminus. The securin^160–202^–separase fusion construct was described previously^11,19^. SCC1-separase fusion constructs were generated by replacing the autocleavage fragment (aa 1482-1536) in insert 2 of separase with SCC1 fragments: aa 100-320 (site 1), aa 310-550 (site 2) or aa 1-631 (full length). In early studies, the C-terminus of SCC1 (aa 1-631) was fused to the N-terminus of separase through a long unstructured linker. SA1/2 subunits were cloned into the pF1 vector with a Twin-StrepII tag at the C-terminus. SMC1A and SMC3 subunits were cloned into the pF1 vector harbouring a dual promoter, with a Twin-StrepII tag at the C-terminus of SMC1A and an 8xHis tag at the C-terminus of SMC3. For fluorescence polarization (FP) experiments, SCC1 (aa 100-320) and SCC1 (aa 350-550) were cloned into a pETM41 vector containing an N-terminal 6xHis-MBP tag and a C-terminal tetracysteine FCM motif (FLNCCPGCCMEP) for fluorescein arsenical hairpin (FlAsH) labelling^39^. The longer SCC1 fragment (aa 100-550) was cloned into the pETM41 vector with a His-MBP tag at the N-terminus. The kinase domain of the human polo-like kinase PLK1 (aa 37-338) in pETM41 vector was kindly provided by Claudio Alfieri (ICR, London). All deletions and mutations were generated by mutagenesis PCR.

### Expression and protein purification

Human separase constructs and cohesin subunits were expressed with a baculovirus expression system. Typically, 25 ml of recombinant P3 baculoviruses were used to infect 500 ml of Sf9 insect cells (Invitrogen) at a cell density of 2.0×10^6^ to 3.0×10^6^ cells/ml. Infected cells were incubated at 27 °C with shaking at 100 rpm for approximately 48 h. Cells were harvested at a cell viability rate of 80-90%, flash-frozen in liquid nitrogen and stored at –80 °C until further use. SCC1 constructs (aa 100-320, 350-550 and 100-550) and PLK1 (aa 37-338) were expressed in *Escherichia coli* BL21 (DE3) cells at 18 °C for 16 h after induction with 0.5 mM IPTG (Isopropyl-β-D-1-thiogalactopyranoside).

Purification of all proteins and protein complexes was performed at 4 °C. Cell pellets of 4 l Sf9 cultures expressing apo-separase or SCC1-separase fusion complexes were resuspended in lysis buffer (50 mM HEPES-KOH pH 8.0, 500 mM KCl, 5% glycerol, 0.5 mM EDTA, 0.5 mM TCEP) supplemented with protease inhibitor cocktail tablets (PIC) (Complete EDTA-free; Roche Diagnostics GmbH), and 5 units/ml SuperNuclease (Sino Biological). The resuspended cells were lysed by sonication and centrifuged at 18,000 rpm for 1 h. The clarified lysate was slowly (0.8 ml/min flow rate) loaded onto two tandem 5 ml StrepTactin Superflow Cartridge (Qiagen) and the columns were washed with the lysis buffer until stable UV absorption was observed. Proteins were eluted in lysis buffer containing 2.5 mM D-desthiobiotin. The eluate was diluted with salt-free buffer to a final KCl concentration of 150 mM and loaded onto a 5 ml Hitrap Heparin HP column (Cytiva) equilibrated in a buffer of 50 mM HEPES-KOH pH 8.0, 150 mM KCl, 5% glycerol, 0.5 mM EDTA, 0.5 mM TCEP. Separase proteins were eluted using a linear gradient of KCl from 150 mM to 1 M. Peak fractions from the Heparin column were pooled, concentrated and passed through a Superose 6 Increase 10/300 GL column (GE Healthcare Life Sciences), pre-equilibrated in gel filtration buffer (20 mM HEPES-KOH pH 8.0, 300 mM KCl, and 0.5 mM TCEP). The purified proteins were pooled, concentrated, flash-frozen and stored at –80 °C for further use. Purification of securin^160–202^–separase fusion protein was performed as described previously^11,19^ and the protein was stored in a buffer containing 20 mM HEPES-KOH pH 8.0, 100 mM KCl, 10 mM MgCl_2_, 0.5 mM TCEP and 5% glycerol.

SA1 and SA2 were purified using StrepTactin Superflow Cartridge (Qiagen) columns. The lysis and washing buffer consisted of 50 mM HEPES-KOH pH 8.0, 500 mM KCl, 5% glycerol and 0.5 mM TCEP. The columns were extensively washed until a stable UV absorption was reached. SA1/2 proteins were eluted using wash buffer supplemented with 2.5 mM D-desthiobiotin. Fractions containing SA1/2 were concentrated and loaded on a Superose 6 Increase 10/300 GL column (GE Healthcare Life Sciences). The purified SA1/2 proteins were stored in gel filtration buffer (20 mM HEPES-KOH pH 8.0, 300 mM KCl and 0.5 mM TCEP) at –80 °C. For expression of SCC1^310–550^–SA2-separase^C2029S/ΔAIL1^ complex, recombinant P3 baculoviruses of SCC1^310–550^–separase^C2029S/ΔAIL1^ fusion construct and SA2 were used to co-infect 3 l of Sf9 cells. The SCC1^310–550^–SA2-separase^C2029S/ΔAIL1^ complex was purified as described for SA1/2 proteins.

The cohesin-separase fusion complex was expressed in Sf9 cells by co-infection with recombinant P3 baculoviruses of SMC1A-SMC3 subunits, SCC1^1–631^–separase^C2029S/ΔAIL1^ fusion construct and SA2 subunit. Pellets of 5 l Sf9 cells were lysed by sonication in lysis buffer (50 mM HEPES-KOH pH 8.0, 300 mM KCl, 5% glycerol, 0.5 mM EDTA, and 0.5 mM TCEP) supplemented with protease inhibitors and SuperNuclease. The lysate was centrifuged at 18,000 rpm for 1 h. The supernatant was loaded onto StrepTactin Superflow Cartridge (Qiagen) columns at a flow rate of 0.8 ml/min and proteins were eluted in lysis buffer containing 2.5 mM D-desthiobiotin. KCl concentration of the eluate was diluted to 150 mM and the complex was further purified using a 5 ml Hitrap Heparin HP column (Cytiva). Peak fractions containing cohesin-separase fusion complexes were pooled, concentrated and loaded on a Superose 6 Increase 10/300 GL column (GE Healthcare Life Sciences), pre-equilibrated in gel filtration buffer (20 mM HEPES-KOH pH 8.0, 300 mM KCl, and 0.5 mM TCEP). The purified proteins were concentrated to approximately 3.2 mg/ml, flash-frozen and stored at –80 °C.

MBP-tagged SCC1 fragments (aa 100-320, 350-550, and 100-550) were purified using a Hitrap MBP column. Briefly, *E. coli* cells expressing SCC1 fragments were resuspended in lysis buffer (25 mM HEPES pH 7.7, 500 mM NaCl, 1 mM EDTA, 2 mM DTT). After centrifugation, the supernatant was filtered and applied to a 5 ml Hitrap MBP column equilibrated in lysis buffer. Proteins were eluted using a buffer containing 25 mM HEPES pH 7.7, 150 mM NaCl and 2 mM DTT. Fractions containing SCC1 were pooled, concentrated to 1 ml and loaded on a Superdex 200 Increase 10/300 GL gel filtration column (GE Healthcare Life Sciences) pre-equilibrated in 25 mM HEPES pH 7.7, 150 mM NaCl and 2 mM DTT. Purification of PLK1 was performed as previously described^28^.

### Fluorescence anisotropy

SCC1 site 1 (aa 100-320) and SCC1 site 2 (aa 350-550) were labelled with the FlAsH-EDT2 fluorescent dye using a buffer containing 25 mM HEPES pH 8.0, 150 mM NaCl and 1 mM β-mercaptoethanol. The labelling reaction was initiated by mixing 5 µM protein with 25 µM dye, followed by an overnight incubation at 4°C. The excess dye in the samples was removed by gel filtration.

Fluorescent anisotropy assays were performed to investigate the molecular interactions between separase and SCC1 (aa 100-320) or SCC1 (aa 350-550). All measurements were conducted using a fluorescence spectrometer equipped with polarizers in both the excitation and emission pathways. The assays were performed in buffer containing 25 mM HEPES pH 8.0, 100 mM NaCl, 0.1 mg/ml BSA and 0.01% NP40. For each experiment, the fluorescently labelled probe was excited at BLA, and the emitted light was measured at BLA. The fluorescence intensity was recorded in parallel (ΔI∥⊥) and perpendicular (ΔI⊥) planes relative to the polarization of the excitation beam. These intensities were used to calculate the anisotropy (r) according to the equation:

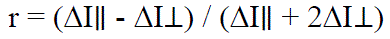

Each measurement was performed at room temperature (25°C). Data were normalized using GraphPad Prism, with additional fitting performed using the One Site-Total non-linear fit equation.

### Separase cleavage assays

^35^S-methionine-labeled human SCC1 was produced in vitro using the TnT® coupled wheat extract systems (Promega) according to the manufacturer’s instructions. The system utilized the TnT® T7 Wheat Germ Polymerase, with 0.8 μCi/μl of ^35^S-methionine added to a 50 μl reaction mix, followed by incubation for 1 h at 30°C. For phosphorylation of SCC1, 4 μM PLK1 was incubated with SCC1 in a buffer containing 25 mM HEPES pH 7.5, 5 mM ATP, 10 mM MgCl₂, and 2 mM DTT for 2 h at 30°C. The cleavage reaction was initiated by mixing 0.25 µM securin^160–202^–separase fusion protein with SCC1 substrate in a buffer containing 25 mM HEPES pH 7.7, 5 mM MgCl_2_, 1 mM DTT and KCl at 30°C. The reaction was terminated by adding 3x SDS-loading buffer. Reaction products were separated by SDS-PAGE using a 4–20% gel (Bio-Rad) and visualized with a Phosphorimager (Typhoon™ FLA 9500). Quantification of band intensities was performed using ImageJ software^40^.

### Phosphorylation of SCC1-separase fusion complexes

Purified SCC1^100–320^–separase^C2029S/ΔAIL1^ or SCC1^310–550^–separase^C2029S/ΔAIL1^ fusion complex was mixed with the kinase domain of PLK1 at a molar ratio of 1:2 in a reaction buffer containing 50 mM HEPES pH 8.0, 150 mM NaCl, 10 mM MgCl_2_, 5 mM ATP, 0.5 mM TCEP and 5% glycerol. The reaction mixture was incubated at room temperature for 1 h and subsequently applied to gel filtration on a Superose 6 increase 5/150 column (GE Healthcare Life Sciences) equilibrated with 20 mM HEPES pH 8.0, 100 mM KCl, and 0.5 mM TCEP. Fractions containing SCC1-separase fusion complexes was directly used for cryoEM grid preparation.

### Mass spectrometry analysis of phosphorylated SCC1 (aa 100-550)

Purified SCC1 (aa 100-550) was incubated with 18 µg PKL1 for 1 h at 30 °C in a buffer containing 50 mM HEPES pH 8.0, 150 mM NaCl, 10 mM MgCl_2_, 5 mM ATP, 0.5 mM TCEP and 5% glycerol. The reaction mixture was subjected to gel filtration to remove PLK1. Peak fractions containing SCC1 (aa 100-550) were pooled and used for mass spectrometry analysis of phosphorylation sites.

### Negative staining analysis of cohesin-separase incubated with DNA

DNA sequence from the pericentromeric region of the centromere on human chromosome 18 was cloned into a high-copy plasmid containing two flanking EcoRV restriction sites. Plasmids were amplified in DH5a cells for 16-18 h. The 340-bp DNA was purified as previously described^41^. Final purification of the DNA was performed by anion exchange chromatography. Purified DNA was stored in a buffer containing 20 mM HEPES pH 8.0, 200 mM KCl, and 1 mM EDTA for further use.

Freshly purified cohesin-separase^C2029S/ΔAIL1^ fusion complex was mixed with 340-bp double-stranded DNA in a molar ratio of 1:1.5 and incubated for 30 min on ice. The sample was then incubated with 1 mM bis(sulfosuccinimidyl)suberate (BS3) in a buffer of 20 mM HEPES pH 8.0, 100 mM KCl, and 0.5 mM TCEP for another 30 min on ice to allow crosslinking. The reaction was terminated by adding 30 mM Tris-HCl pH 8.0. 3 µl cross-linked sample was applied to carbon-coated 400-mesh copper grids (Electron Microscopy Sciences) that had been glow discharged for 25 s. The sample was incubated on the grid for 1 min before blotting with filter paper to remove excess liquid. Grids were washed with water and stained with 2% (w/v) uranyl formate. A total of 3,508 micrographs were acquired on a Talos L120C microscope operated at 120 kV with a nominal magnification of 57,000x, resulting in a pixel size of 2.463 Å. Images were recorded using EPU software with a defocus range of –1.5 to –2.0 µm. Image processing was performed in CryoSPARC v.4.4.1^42^. 302,050 particles of separase bound to SMC subunits were selected and subjected to 3D classification. Three distinct classes showed extra density bound to separase. Full length human separase, and AF3 predicted structures of SMC1-SCC1 and SMC3-SCC1 complexes were fitted into the negative staining EM density maps.

### CryoEM sample preparation and data collection

Purified apo-separase (active and inactive) was diluted in a buffer of 20 mM HEPES pH 8.0, 100 mM KCl, and 0.5 mM TCEP, and then applied to graphene oxide-coated holey-carbon grids (Quantifoil R1.2/1.3, 300 mesh, gold) at a concentration of 150-200 nM. Graphene oxide grids were prepared following the protocol described in Boland *et al*.^29,43^. The grids were incubated for 10 s and back blotted for 2 to 3 s with 1 mm additional movement (90% humidity at 16°C), and then plunged into liquid ethane using a Leica EM GP2 automatic plunge freezer. Freshly prepared phosphorylated SCC1-separase fusion complexes were dispensed onto graphene oxide-covered Quantifoil R1.2/1.3 holey-carbon grids at a concentration of 100 to 200 nM. The prepared grids were directly screened and subsequently used for cryoEM data collection. All data sets were acquired on a Thermo Scientific Talos Arctica Cryo-TEM operating at an accelerating voltage of 200 kV, equipped with either Falcon 3 or Falcon 4i direct electron detector. Data acquisition was monitored on-the-fly pre-processing using CryoSPARC v.4.4.1.

For active apo-separase a total of 3,314 movies and for inactive apo-separase^C2029S^ a total of 5,292 movies were recorded using a Falcon 3 direct electron detector at a nominal magnification of 150,000x, resulting in a pixel size of 0.9759 Å. Data were collected using EPU (Thermo Fisher Scientific) with one image per hole at a defocus range of –0.6 to –2.2 μm and a total electron dose of 40 e^-^/Å^2^ distributed over 40 frames per acquisition.

For phosphorylated SCC1^100–320^–separase^C2029S/ΔAIL1^ or SCC1^310–550^–separase^C2029S/ΔAIL1^ fusion complexes, a total of 12,867 movies or 7,451 movies were recorded, respectively, using a Falcon 4i direct electron detector at a nominal magnification of 130,000x, resulting in a pixel size of 0.9024 Å. A post-column energy filter (Selectris X) was used for zero-loss filtration with an energy width of 10 eV. Data were collected using EPU (Thermo Fisher Scientific) with one image per hole, a defocus range of –0.6 to –2.0 μm and a total electron dose of 50 e^-^/Å^2^ distributed over 50 frames per acquisition.

Purified SCC1^310–550^–SA2-separase ^C2029S/ΔAIL1^ complex was diluted to a final concentration of either 0.3 mg/ml or 150-200 nM in a buffer of 20 mM HEPES pH 8.0, 150 mM KCl, and 0.5 mM TCEP. 5 µl of diluted sample (at 0.3 mg/ml) was applied onto holey-carbon grids (Quantifoil R1.2/1.3, 300 mesh, gold), front blotted for 3-3.5 s with 1 mm additional movement (95% humidity at 15°C) before being plunged into liquid ethane using a Leica EM GP2 automatic plunge freezer. Alternatively, 5 µl of diluted sample (at 150-200 nM) was applied onto graphene oxide-coated grids, back blotted for 2-2.5 s with 1 mm additional movement (95% humidity at 15 °C) and then plunged into liquid ethane using the same plunge freezer. A total of 13,810 movies and 31,617 movies were recorded using a Falcon 4i direct electron detector on standard grids and graphene oxide-coated grids, respectively. Data acquisition was performed at a nominal magnification of 130,000x, resulting in a pixel size of 0.9024 Å, with an energy filter slit width of 10 eV. Each data set was collected with a total electron dose of 50 e^-^/Å^2^ distributed over 50 frames. Data were collected using EPU (Thermo Fisher Scientific) with 3 images per hole and a defocus range of –0.6 to –2.0 μm. To attenuate the preferred orientation of particles, additional 11,585 movies were recorded on graphene oxide-coated grids with a stage tilt of 30° at the same magnification. Data collection for the tilted data set was performed with a set defocus range of –1.0 to –2.6 μm.

For cryoEM grid preparation of cohesin-separase ^C2029S/ΔAIL1^-DNA complex, 4 µl of cross-linked sample was applied onto ANTcryo grids (R1.2/1.3, 300 mesh, gold) with an amorphous nickel titanium alloy film. The grids were front blotted for 3–3.5 s with 1 mm additional movement under 95% humidity at 15°C and then plunged into liquid ethane. A total of 22,985 movies were recorded using a Falcon 4i direct electron detector at a nominal magnification of 130,000x, resulting in a pixel size of 0.9024 Å. Data collection was performed using EPU (Thermo Fisher Scientific) with three images per hole, a defocus range of –0.8 to –2.4 μm and a total electron dose of 50 e^-^/Å^2^ distributed over 50 frames.

### CryoEM image processing

All data were processed using CryoSPARC v.4.4.1^42^ and RELION 4.0^44^. The data processing workflows are summarized in the Extended Data Figures. Raw movies for all data sets were aligned and dose-weighed using patch-based motion correction, and contrast transfer function (CTF) parameters were estimated by patch-based CTF estimation in CryoSPARC.

For the wild-type apo-separase data set, particles were initially picked using a blob picker with a minimum and maximum diameter of 120 Å and 220 Å. The resulting 2D class averages were used as templates for template-based picking. Particle clean-up using three rounds of 2D classification resulted in 654,549 particles followed by two rounds of 3D classification. 224,027 particles were selected and subjected to global CTF refinement, local motion correction and non-uniform refinement, resulting in a map that refined to 3.3 Å resolution. To improve the local density of separase, local refinement was performed using masks focused on the HEAT-repeat domain or the TPR-like and protease domains (SPD). A composite map was generated by combining two focused refinement maps.

For the apo-separase^C2029S^ data set, initial particles were picked by template picker using the templates generated from the active wild-type apo-separase reconstruction. After particle clean-up through three rounds of 2D classification, 852,228 particles were retained for subsequent 3D classification. Resulting 368,728 particles were subjected to global CTF refinement, local motion correction and non-uniform refinement, resulting in a map at 3.2 Å resolution. Particles were further cleaned up by another round of 2D classification, followed by 3D refinement generating a map at 3.15 Å resolution. To improve the density of separase autocleavage sites, 3D classification without alignment was performed using a mask covering TPR-like domain and SPD. Four classes of particles (118,288 particles) showing improved density of autocleavage sites were selected for 3D refinement, yielding a map at 3.1 Å resolution. Local resolution of TPR-like domain and SPD was further improved by local refinement.

For the phosphorylated SCC1^100–320^–separase^C2029S/ΔAIL1^ data set, templates generated from apo-separase^C2029S^ reconstruction were used and subsequently cleaned up by four rounds of 2D classification and two rounds of 3D classification. 518,130 particles were used for non-uniform refinement and CTF refinement and resulted in a reconstruction of 2.9 Å resolution. To improve the occupancy of SCC1, refined particles were imported into RELION for 3D classification without alignment (K=5 and T=8), using a mask on SCC1. The class (214,707 particles) showing the strongest SCC1 density was selected, and particles were subjected to CTF refinement and non-uniform refinement. Another round of 3D classification without alignment (K=6 and T=12) was performed in RELION, using a mask covering the TPR-like and protease domains and SCC1. The best class showing high-resolution features was selected (195,231 particles) for 3D refinement in CryoSPARC, resulting in a map at 2.9 Å resolution. Local refinement was performed to further improve the density of SCC1.

For the phosphorylated SCC1^310–550^–separase^C2029S/ΔAIL1^ complex, the processing strategies were like those described above. Briefly, after data clean-up through 2D and 3D classifications, 298,574 particles were retained for subsequent CTF refinement and non-uniform refinement, followed by local refinement focused on the TPR-like and protease domains. This resulted in a reconstruction at a resolution of 2.9 Å. 3D classification without alignment (K=6 and T=8) in RELION using a mask on the HEAT-repeat domain of separase resulted in a class (180,532 particles) with clear density of SCC1 binding to separase. The particles were re-imported into CryoSPARC for non-uniform refinement followed by a final local refinement with a soft mask around HEAT-repeat domain. As a result, the density of SCC1 binding to HEAT-repeat domain was significantly improved.

For the SCC1^310–550^–SA2-separase^C2029S/ΔAIL1^ complex, untilted and tilted (30°) data sets were processed separately until the 2D classification step. Initial particles were picked by a template picker combined with a blob picker and cleaned up through several rounds of 2D classification. Particles corresponding to the tertiary complex, SCC1^310–550^–separase and SCC1-SA2 complexes from each data set were selected and merged for further processing. A total of 795,457 particles for the tertiary complex were subjected to four rounds of 3D classification. The resulting 382,898 particles were used for CTF refinement and non-uniform refinement followed by further data clean-up through 2D and 3D classification. Particles of the best class and particles corresponding to the tertiary complex from other classes were selected (total 177,878 particles). 3D refinement using non-uniform refinement generated a reconstruction at 3.4 Å resolution. Local refinement using a soft mask 1 covering SCC1, separase and C-terminal helices of SA2 significantly improved the density of the interaction interface and reduced the anisotropy of 3D reconstruction, producing a map at 3.1 Å resolution. A soft mask 2 on SA2, SCC1 and part of the protease domain was also applied to local refinement improving the map of SCC1-SA2. A composite map of the tertiary complex was created by combining two focused-refined maps. 3D classification of 1,375,401 particles for SCC1^310–550^–separase complex resulted in a best class with 614,186 particles. Following CTF refinement and non-uniform refinement yielded a map at 2.9 Å resolution. Local refinement using soft masks on each domain further improved the local densities. The focused-refined maps were used to generate a composite map of SCC1^310–550^–separase complex. Similarly, after 3D classification of total 2,823,666 particles for SCC1^310–550^–SA2 complex, the best class containing 768,540 particles was selected for further CTF refinement and non-uniform refinement. Subsequent local refinement was performed using soft masks covering the N-terminal part (mask 3) or C-terminal part (mask 4) of SA2. The two locally improved maps were combined to produce a composite map.

For the cohesin-separase ^C2029S/ΔAIL1^-DNA complex data set, particles were initially picked using both blob template pickers. Following several rounds of 2D classification and removal of duplicated particles, resulting 2D class averages showing separase bound to cohesin subunits were used as input for the Topaz particle-picking pipeline to increase the number and accuracy of picked particles. After clean-up through multiple rounds of 2D and 3D classifications, 179,047 particles were subjected to CTF refinement and non-uniform refinement. The resulting map refined to an overall resolution of 3.4 Å. To improve the density of cohesin subunits bound to separase, 3D classification without alignment (six classes) was performed in CryoSPARC. One class (26,747 particles) showing enhanced density of cohesin subunits was selected for a final non-uniform refinement, producing a reconstruction of 4.3 Å. All maps were post-processed with a three-dimensional deep learning framework named EMReady^45^. All resolution estimations were derived from Fourier shell correlation (FSC) calculations between reconstructions from two independently refined half-maps. Reported resolutions are based on the FSC gold standard criterion. Local resolution estimations are obtained by ResMap^46^.

### Model building and refinement

The cryoEM structure of human separase-securin complex (7NJ1) was used as an initial reference to model the structures of apo-separase (active and inactive), SCC1^100–320^–separase^C2029S/ΔAIL1^ and SCC1^310– 550^–separase^C2029S/ΔAIL1^ fusion complexes. Model building of SCC1 in regions that were difficult to interpret was facilitated by automated atomic model building using ModelAngelo^47^ and Alphafold predictions^31^. Similarly, the crystal structure of SCC1-SA2 (4PJU^36^) together with the SCC1^310–550^– separase^C2029S/ΔAIL1^ structure from this study were used as initial models for building the structure of the SCC1^310–550^–SA2-separase^C2029S/ΔAIL1^ complex. All initial models were fitted into cryoEM maps using Chimera X^48^, manually built in COOT^49^, and real-space refined using PHENIX^50^. Model validation was performed in MolProbity^51^. Structural figures were generated in Chimera X. All the statistics are summarized in **Extended Data Table 1**.

### Sample preparation for Crosslinking Mass spectrometry

The purified cohesin-separase^C2029S/ΔAIL1^ fusion complex was mixed with 340-bp double-stranded DNA in a molar ratio of 1:1.2 and incubated for 30 min on ice. The sample was then incubated with 0.25 mM sulfosuccinimidyl 4,4’-azipentanoate (Sulfo-SDA, Thermo Scientific Pierce) in a buffer of 20 mM HEPES pH 8.0, 100 mM KCl, and 0.5 mM TCEP for 10 min at room temperature. The mixture was transferred onto Eppendorf tube lids, placed on ice and irradiated with UV light at 365 nm for 10 seconds using a LED Spot 100 HP IC (honle uv technology). Reactions were quenched with 50 mM ammonium bicarbonate for 10 minutes and then precipitated by adding four volumes of −20 °C cold acetone and incubating for overnight at −20 °C. Precipitated material was pelleted by centrifugation at 20,000g for 15 min. The pellet was briefly washed in cold acetone, pelleted again and air-dried. The crosslinked samples were resuspended in 8 M urea, 100 mM ammonium bicarbonate. Proteins were reduced with 5 mM DTT and alkylated with 10mM iodoacetamide and digested with Lysyl Endopeptidase (Lys-C) at an enzyme-to-protein ratio of 1:100 for 4 h at room temperature with shaking. Urea was diluted to 1.5 M with 100 mM ammonium bicarbonate solution and the peptides were further digested with trypsin at an enzyme-to-protein ratio of 1:50 overnight at room temperature with shaking.

Digested peptides were cleaned and desalted by C18 stagetips and were dried in a vacuum concentrator (Eppendorf). For crosslinked peptide enrichment, peptides were fractionated on an ÄKTA Pure system (GE Healthcare) using a Superdex 30 Increase 3.2/300 (GE Healthcare) at a flow rate of 10 μl per minute using 30% (v/v) acetonitrile and 0.1% (v/v) trifluoroacetic acid as the mobile phase at 4°C. Fifty-microlitre fractions were collected from the elution volume 1.00 ml to 1.45 ml and dried for subsequent liquid chromatography–tandem mass spectrometry (LC–MS/MS) analysis.

### Liquid chromatography and cross-linking mass spectrometry (LC-XLMS)

All peptides were resuspended in 0.1% formic acid (FA) and 1.6% acetonitrile (ACN) before being analyzed on a Thermo Eclipse Orbitrap MS coupled to a Vanquish Neo HPLC, using an EASY-Spray source and PepMap Neo 75 µm × 500 mm C18 column at 0.3 µl/min. A 95-minute gradient was employed (1.6% to 44% ACN/0.1% FA), followed by 44% to 76% ACN/0.1% FA in 2.5 minutes. Global MS parameters included a 95-minute total method, advanced peak determination checked, default charge state of 2, EASY-IC internal calibration, static spray voltage of 2 kV, sweep gas of 2, ITT temperature 280 °C, expected LC peak widths of ∼30 s, and positive polarity. Fractions 1–9 were analyzed with different acquisition methods (frac1:A, frac2:A/B/E, frac3:A/B/C/E, frac4-9: A/D/E). Acquisition A (duty cycle 3 s) used an Orbitrap MS1 at 240,000 resolution (m/z 380–2,000), quadrupole isolation, AGC target of 150% (max injection time 100 ms), one microscan in profile mode, and dynamic exclusion (±10 ppm, 10 s) for charge states 3–7, with subbranches defining precursor selection ranges for 4+ (m/z 380–1,800), 5+ (m/z 380–1,350), and 6–7+ (m/z 380–1,000). ddMS2 scans were Orbitrap HCD at 60,000 resolution (m/z 150–2,000), isolation window of 1.4 m/z, stepped collision energies of 20, 26, and 30 (normalized), an AGC target of 750% (max injection time 150 ms), one microscan in centroid mode, and charge 3+ as second priority. Acquisition B, C, and D followed the same setup but with targeted precursor selection ranges: B (m/z 750–1,005), C (m/z 1,000–1,800), and D with three windows (m/z 380–655, 650–905, and 900–1,800, respectively). Acquisition E (“Boxcar”) used a 5.12 s duty cycle with targeted SIM at 240,000 resolution, multiplex isolation of 10 ions (user-defined groups), custom AGC target with auto injection time, and source fragmentation at 10 V; two boxcar mass lists specified the center m/z values, isolation windows [Box1: 414.1/30.2, 462.2/24.8, 504.1/23.6, 545.35/23.1, 587.8/25, 634.05/26.7, 686/31.4, 784.45/38.9, 829.65/48.5, 957.2/91.6], [Box2: 439.5/26.6, 483.45/23.7, 524.85/23.9, 566.15/24.3, 610.5/26.4, 658.85/28.9, 715.35/33.3, 786.65/43.5, 882.65/63.5, 1101/202] and an AGC target of 50%. All other parameters were consistent with Acquisitions A–D.

### Data analysis of crosslinking MS

A recalibration to control for detector error was conducted on MS1 and MS2 based on the median mass-shift of high-confidence (<1% FDR) linear peptide identifications from each raw file. To identify cross-linked peptides, the recalibrated peak lists were searched against the sequences and the reversed sequences (as decoys) of cross-linked peptides using the Xi software suite (v.1.8.6; https://github.com/Rappsilber-Laboratory/XiSearch). The following parameters were applied for the search: MS1 accuracy = 2 ppm; MS2 accuracy = 5 ppm; enzyme = trypsin allowing up to 3 missed cleavages and 2 missing monoisotopic peaks; crosslinker = SDA with an assumed NHS-ester reaction specificity for K, Y, S, T and N termini; diazirine reaction specificity for A, Ccm, D, E, G, H, I, K, L, P, S, T, V, Y, C termini and N termini; fixed modifications = carbamidomethylation on cysteine; variable modifications = acetylation on lysine and protein N termini, oxidation on methionine, hydrolysed SDA on lysines and protein N termini. MS cleavage of SDA crosslinks was considered during searches.

Before estimating the FDR, the matches were filtered to those having greater than two fragments matched with a non-cleaved SDA and at least five matches total per peptide. These candidates were then filtered to a 1% FDR at the residue-pair level using XiFDR (v.2.3.2)^52^.

## Acknowledgments

We thank A. Reynaud and Y. Pfister for technical assistance; We thank all group members (Boland and Morgan) for their input and discussion; K. Muir and D. Barford for critical reading of the manuscript. C. Alfieri for sharing his PLK1 plasmid; the computing department of the University of Geneva for providing an infrastructure to perform cryoEM analysis; N. Roggli for maintaining computing in the Molecular Biology department; C. Bauer and S. Barass for their contributions to the cryoEM facility in Geneva (DCI-Geneva aka as cryoGEnic); the Protein Production and Structure Core Facility (PTPSP) at the École Polytechnique Fédérale de Lausanne; the Metabolomics Core Facility at EMBL Heidelberg for mass spectrometry analysis. This work was supported by funding from the Intramural Research Program, National Institutes of Health, National Cancer Institute, Center for Cancer Research (Project ZIA BC 012114) to F.J.O., a grant from the US National Institute of General Medical Sciences (R35-GM118053) to D.O.M, and by funding from the Swiss National Science Foundation (SNSF) (TMSGI3_211581), and the Swiss Cancer Research Foundation (KFS-5453-08-2021) all to A.B.

## Data deposits

The coordinates have been deposited in the PDB under the accession numbers 9HMA, 9HM7, 9HMV, 9HN4, 9HN5, 9HN0, 9HMS and 9HVY. The electron density maps have been deposited in the Electron Microscopy Data Bank under accession numbers EMD-52290, EMD-52288, EMD-52307, EMD-52306, EMD-52297, EMD-52295, EMD-52291, EMD-52294, EMD-52303, EMD-52302, EMD-52300, EMD-52301, EMD-52298, and EMD-52445. The XL-MS dataset used in this study are available in the PRIDE database under accession code PXD061667.

## Author contributions

J.Y. designed, expressed and purified all complexes for cryoEM and functional studies; J.Y., S.S. and M.B. prepared proteins for cleavage and binding assays; S.S., C.M.G. and J.M.G. performed separase cleavage assays; M.B. designed and performed fluorescence polarisation binding assays; K.L. and F.J.O. collected and processed XL-MS data; J.Y. collected EM data with support from A.H. and processed all EM data; J.Y., S.S. and M.B. made the figures. A.B. directed the project and designed experiments together with J.Y. and D.O.M.; J.Y., D.O.M. and A.B. wrote the manuscript with input from all other authors.

## Figure legends

**Extended Data Figure 1.**
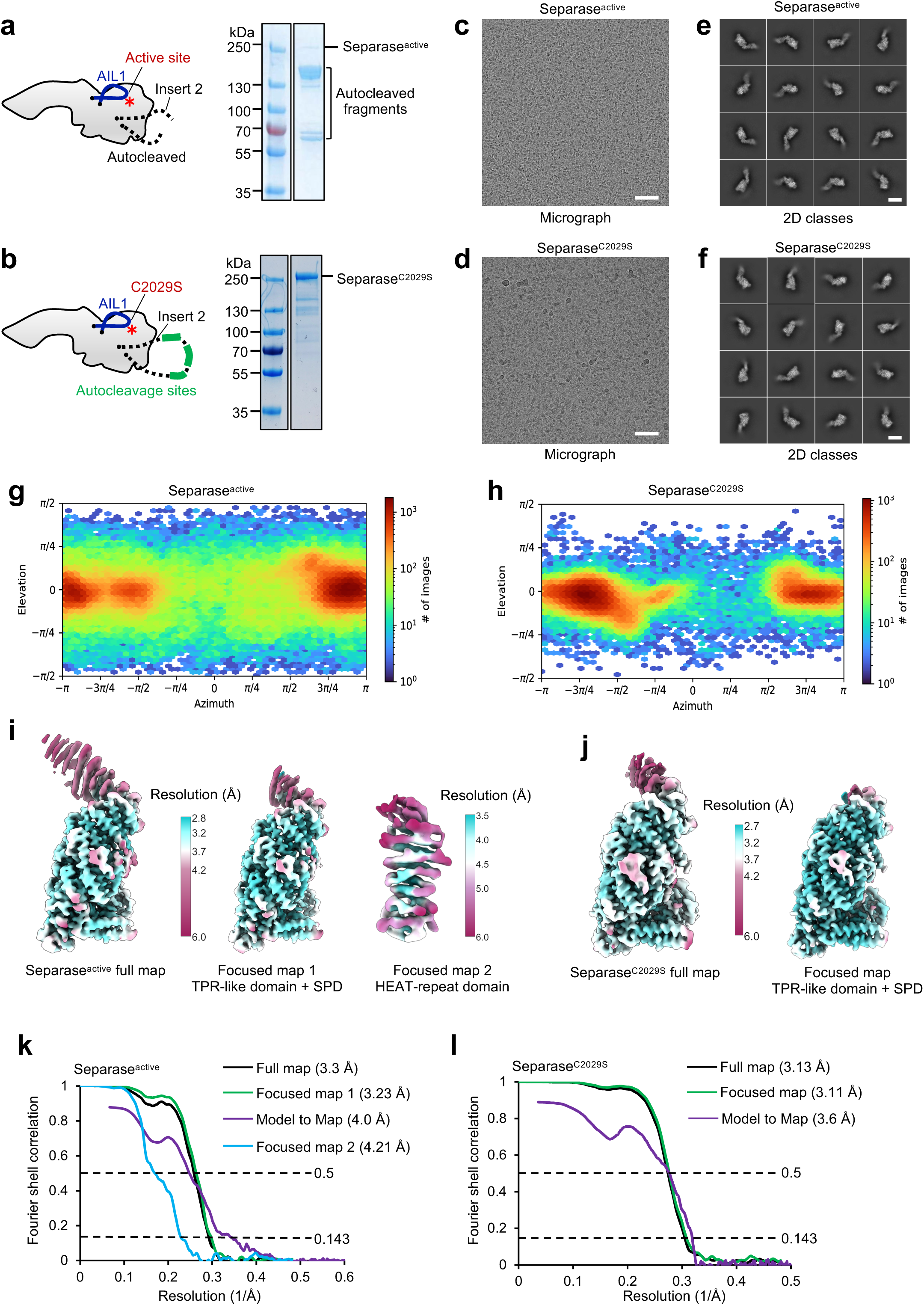
Biochemical and cryoEM analysis of active and inactive apo-separase. **a**, SDS-PAGE gel of active separase, showing that separase undergoes autocleavage during expression and purification. A schematic cartoon that indicates AIL1, cleaved insert 2 and the active site is shown on the left. **b,** SDS-PAGE gel of inactive separase. C2029S mutation prevents separase autocleavage. A schematic cartoon that indicates AIL1, intact insert 2 and the active site is shown on the left. **c-d,** Representative micrographs of active separase (**c**) and inactive separase (**d**), collected on graphene oxide-coated grids to increase the number of views of separase^29,43^. Scale bars, 500 Å. **e-f,** Gallery of two-dimensional class averages of active (**e**) and inactive (**f**) separase, showing typical classes of various views. Scale bars, 100 Å. **g-h,** Angular distribution of active (**g**) and inactive (**h**) separase. Data sets calculated using non-uniform refinement algorithm in CryoSPARC^42^. **i,** EM density maps of active separase colour-coded according to local resolution ranging from 2.8 Å to 6 Å for full map and focussed refined map 1 (TPR-like and protease domains), and 3.5 Å to 6 Å for focussed refined map 1 (HEAT-repeat domain)^46^. **j,** EM density maps of inactive separase colour-coded according to local resolution ranging from 2.8 Å to 6 Å for full map and focussed refined map (TPR-like and protease domains). **k-l,** Gold standard Fourier Shell Correlation (FSC) curves of the full map and the focused refined maps for active separase (**k**) and inactive separase (**l**). The FSC curve between the full cryoEM map and the final atomic coordinates is calculated using Mtriage^53^.

**Extended Data Figure 2.**
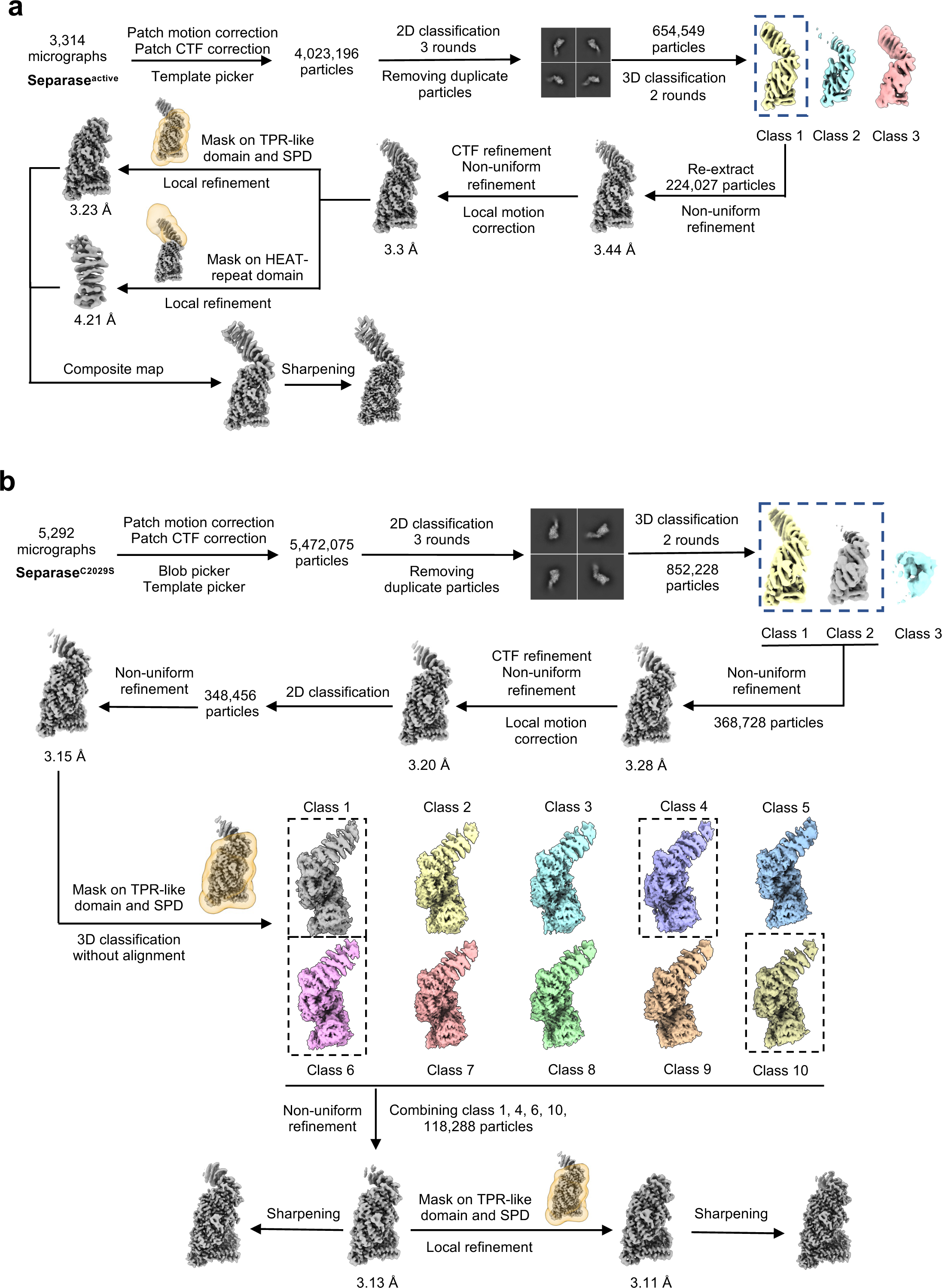
Data-processing flowcharts for both active and inactive apo-separase. **a**, CryoEM processing pipeline of active separase. Initial particles picked from approximately 3,300 micrographs were cleaned up by three rounds of 2D classification and two rounds of 3D classification. The final set of 224,027 particles were subjected to CTF refinement, local motion correction and non-uniform refinement, resulting in a map at 3.3 Å resolution. Local refinement was performed using masks on the TPR-like and protease domains (top), and the HEAT-repeat domain of separase (bottom), yielding maps at 3.2 Å and 4.2 Å, respectively. A composite map was generated by combining the two focussed refined maps. **b,** CryoEM processing pipeline of inactive separase. Following initial cleanup through 2D and 3D classifications, 348,456 particles were selected. These particles were further subjected to non-uniform refinement and 3D classification without alignment, applying a mask on the TPR-like and protease domains. Four classes of particles showing clear density for auto-cleavage fragment were combined and refined, resulting in a final map at 3.1 Å resolution. A final local refinement using a mask on the TPR-like and protease domains was performed. This map also refined to 3.1 Å resolution.

**Extended Data Figure 3.**
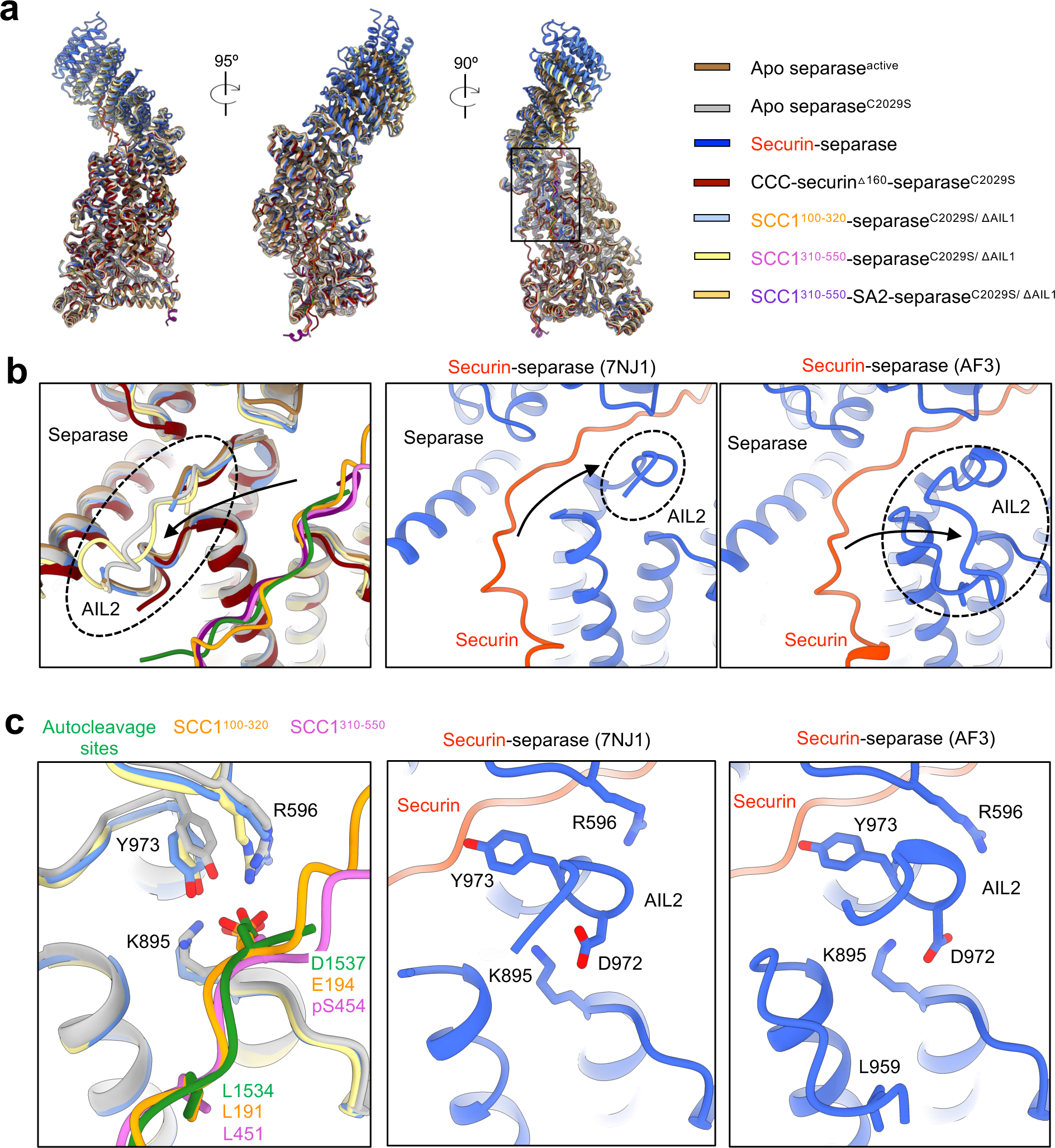
Local conformational changes of separase structures in apo, substrate-bound, and inhibitor-bound states. **a**, Structural alignment of separase bound to substrates, inhibitors and in apo states. Structures are shown as cartoons with separase coloured by different blocks. Securin is in orange red; autocleavage sites are in forest green; SCC1 (aa 100-320) is in orange; SCC1 (aa 310-550) is coloured orchid in the binary complex and purple in the tertiary complex (with SA2). **b,** Conformational change of AIL2. In apo-separase, SCC1-separase and CCC-separase complexes, AIL2 (circled by a dashed line, left panel) binds to a hydrophobic groove on separase. In the securin-separase complex (middle panel), securin occupies the hydrophobic groove while AIL2 binds at the substate-binding site. Alphafold-predicted structure of securin-separase complex reveals the complete AIL2 structure (right panel). The black arrows indicate the movement of AIL2 between complexes. **c,** Close-up view showing AIL2 occupies Lxx[S/D/E] motif-binding site in the securin-separase complex. Left, Lxx[S/D/E] motif from the autocleavage sites and SCC1 site 1 and site 2 binds to the TPR-like domain of separase. Middle and right, residues L959 and D972 of AIL2 occupy Lxx[S/D/E] motif-binding site.

**Extended Data figure 4.**
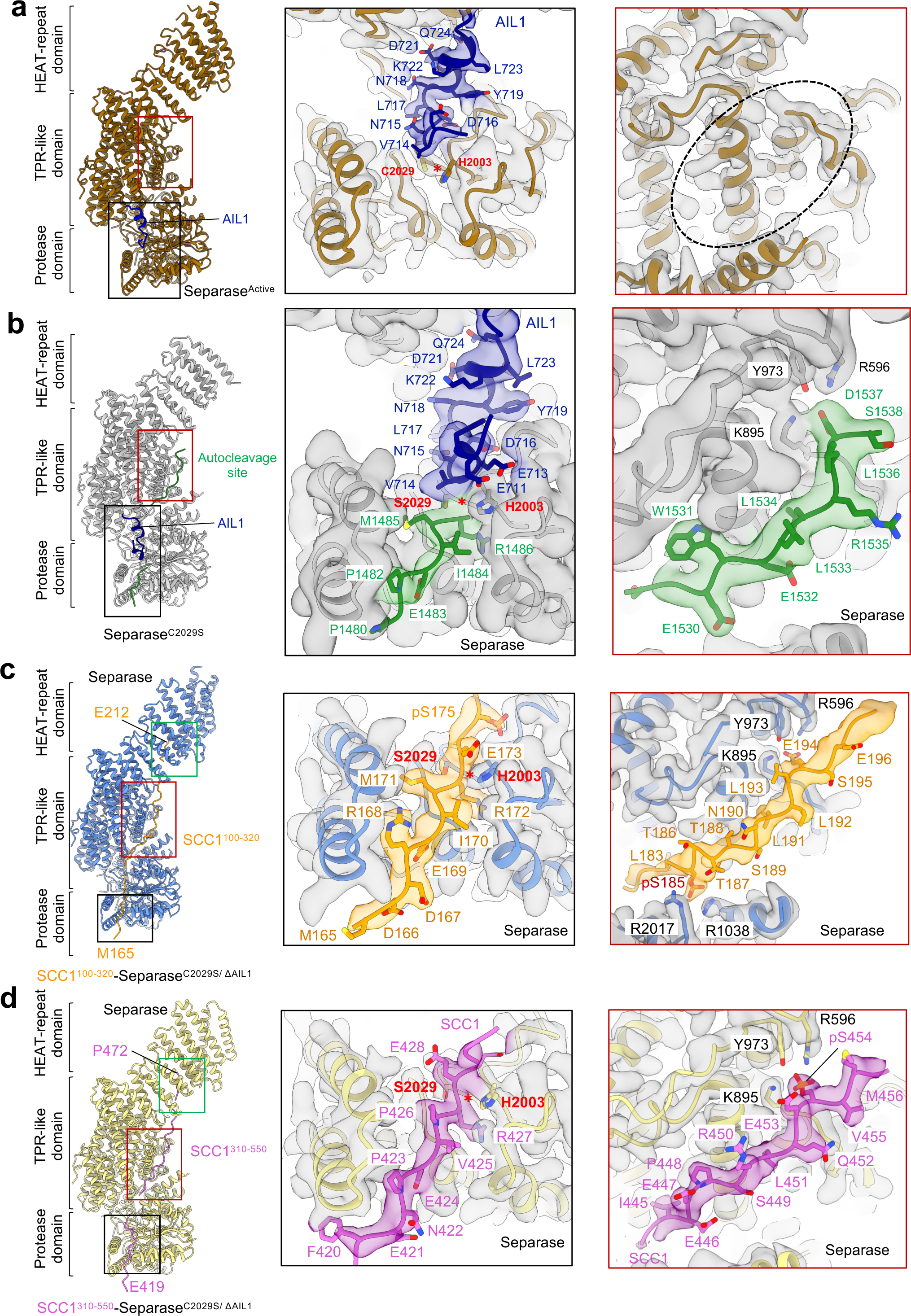
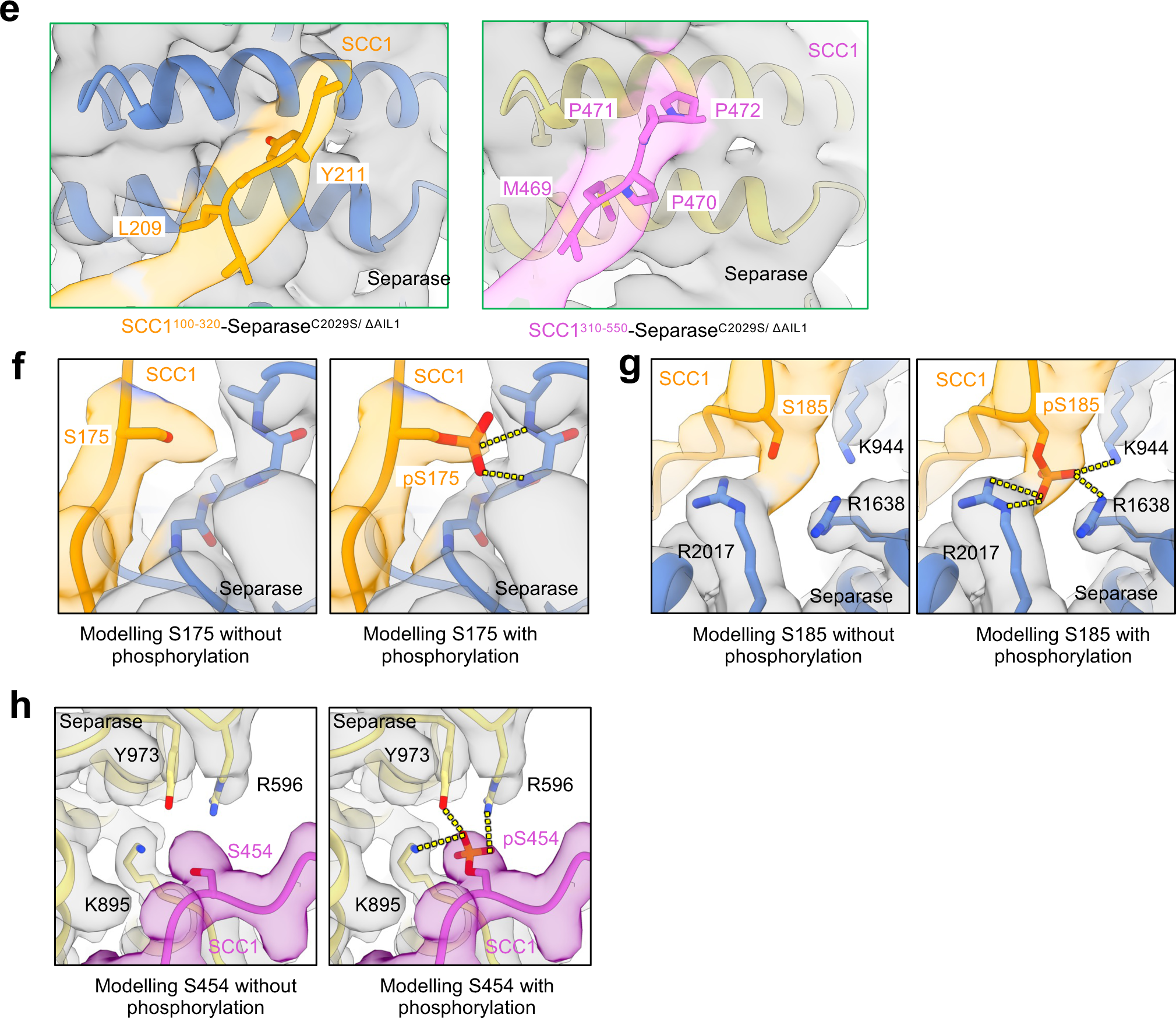
Representative EM density of separase in apo or substrate-bound states. **a**, Ribbon representation of active apo-separase on the left with AIL1 in blue. EM density of AIL1 binding near the catalytic site (black box) and TPR-like domain of active separase (red box). The dashed-line circle indicates no extra density binding in this region. **b,** Ribbon representation of inactive apo-separase on the left with AIL1 in blue and autocleavage sites in green. EM density of the autocleavage site 1 and AIL1 binding to the protease domain (black box), and the autocleavage site 3 binding to the TPR-like domain of inactive separase (red box). **c-d,** Ribbon representation of inactive separase bound to site 1 (**c**) or site 2 (**d**) on the left. EM density of SCC1 (aa 100-320) (**b**) and SCC1 (aa 310-550) (**c**) bound to separase. Black box, density showing the cleavage sites 1 and 2 motifs binding to the protease domain. Red box, density of exosites in SCC1 binding to the TPR-like domain. **e,** Close-up views (green boxes in **b** and **c**) of EM density showing SCC1 site 1 (left) and site 2 (right) bound to the HEAT-repeat domain of separase. **f-h**, EM densities of SCC1 serine residues modelled with and without phosphate group. All densities clearly indicate that S175, S185 and S454 of SCC1 are phosphorylated.

**Extended Data figure 5.**
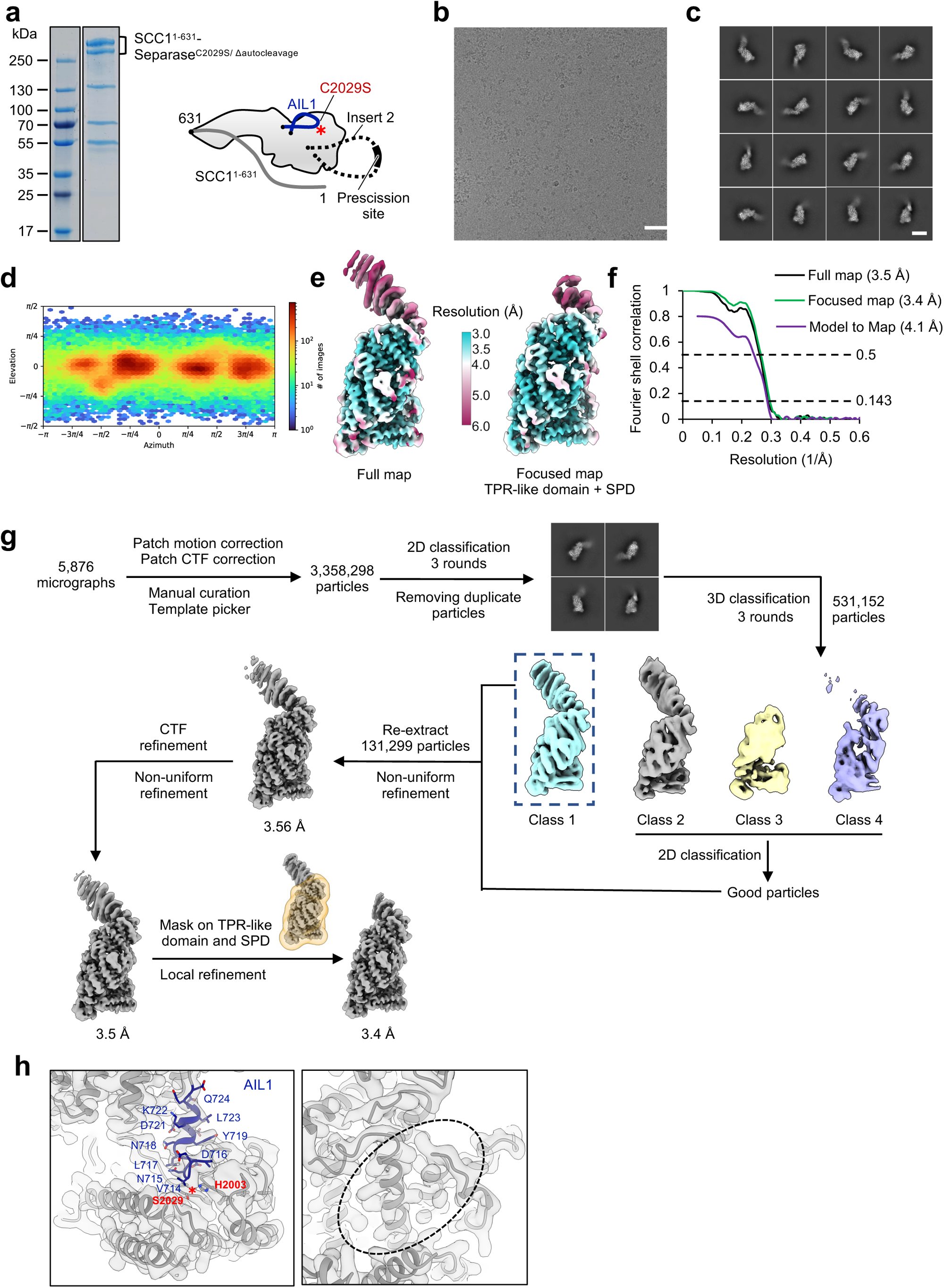
Biochemical and cryoEM analysis of SCC1^1–631^–separase^C2029S/Δautocleavage^ fusion complex. **a**, Schematic representation and SDS-PAGE gel of the fusion complex. In this construct, the C-terminus of SCC1 was fused to the N-terminus of separase with a GS linker in between and the autocleavage sites (aa 1482-1536) within insert 2 was replaced by a preScission site (Δautocleavage). **b,** Representative cryo-electron micrographs of the fusion complex, collected on graphene oxide-coated EM grids to increase the number of views of separase. Scale bars, 500 Å. **c,** Representative two-dimensional class averages of the fusion complex. Scale bars, 100 Å. **d,** Angular distribution plot for the fusion complex data set calculated using non-uniform refinement algorithm in CryoSPARC^42^. **e,** EM density maps of the fusion complex colour-coded according to local resolution. **f,** Gold standard FSC curves of the full map and the focused refined maps. The FSC curve between the full cryoEM map and the final atomic coordinates is calculated using Mtriage^53^. **g,** CryoEM processing pipeline of the fusion complex. The final refinement includes 131,299 particles and the maps refines to a resolution of 3.4 Å. **h,** Representative EM density of AIL1 binding near the catalytic site (left) and the TPR-like domain of separase (right) in the fusion complex. AIL1 is shown in dark blue, and the EM density is shown in light grey. The dashed-line circle indicates no extra density binding in the TPR region as observed for SCC1 site 1 and site 2 or the autocleavage site 3.

**Extended Data figure 6.**
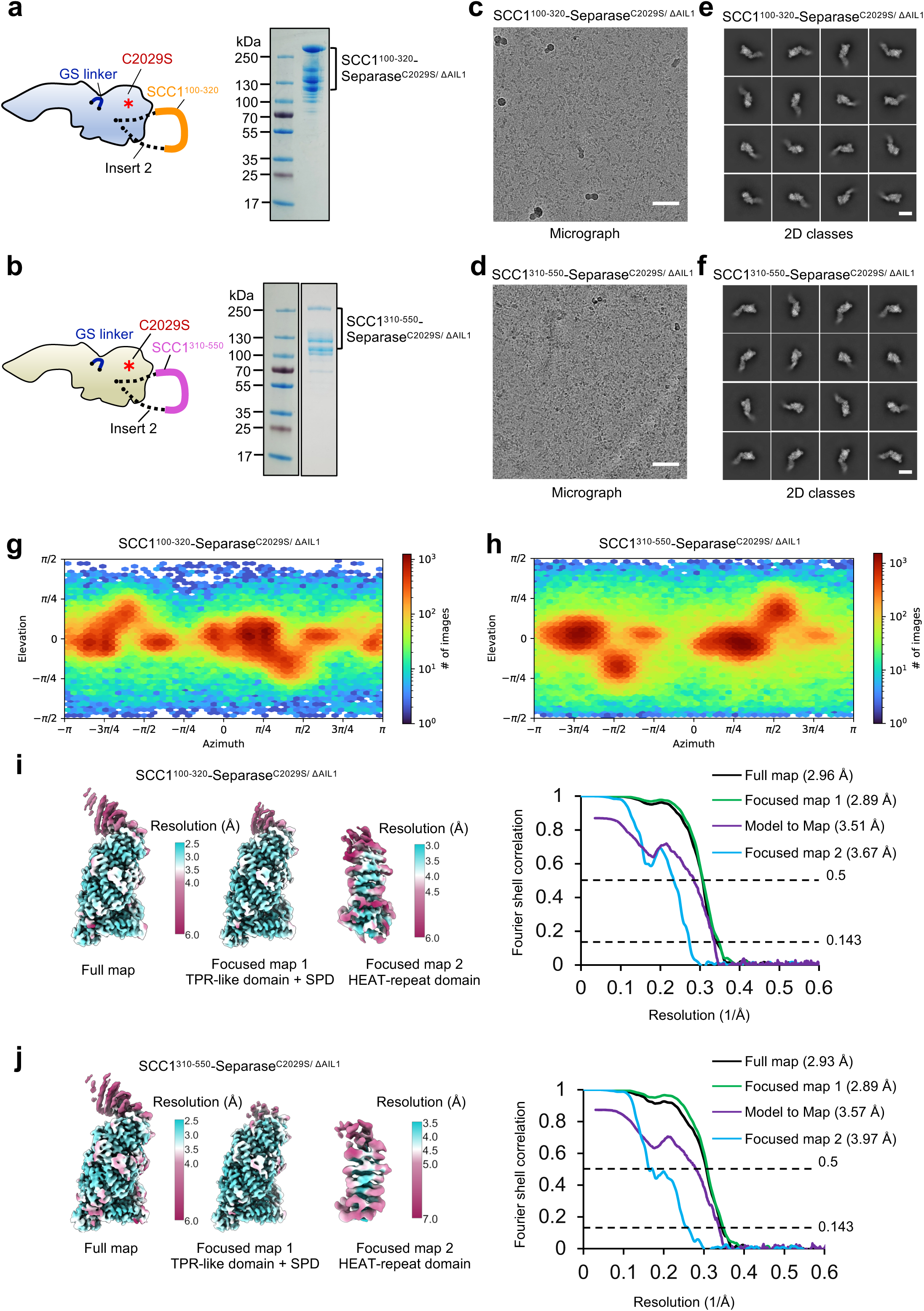
Biochemical and cryoEM analysis of separase bound to SCC1^100–320^ or SCC1^310–550^. **a-b**, SDS-PAGE gels of SCC1^100–320^–separase^C2029S/ΔAIL1^ complex (**a**) and SCC1^310–550^– separase^C2029S/ΔAIL1^ complex (**b**). Here, SCC1 fragments replaced the autocleavage sites (aa 1482-1536) in insert 2 of separase. △AIL1, deletion of AIL1. **c-d,** Representative EM micrographs of the two complexes, collected on graphene oxide-coated grids to increase particle orientation and distribution. Scale bars, 500 Å. **e-f,** Gallery of two-dimensional class averages of the two complexes, showing typical classes of various views. Scale bars, 100 Å. **g-h,** Angular distribution plots for the two complexes calculated using non-uniform refinement algorithm in CryoSPARC^42^. **i-j,** EM density maps of the two complexes colour-coded according to local resolution and gold standard FSC curves for the full map and the focussed refined maps. The FSC curves between the full cryoEM map and the final atomic coordinates were calculated using Mtriage^43^.

**Extended Data figure 7.**
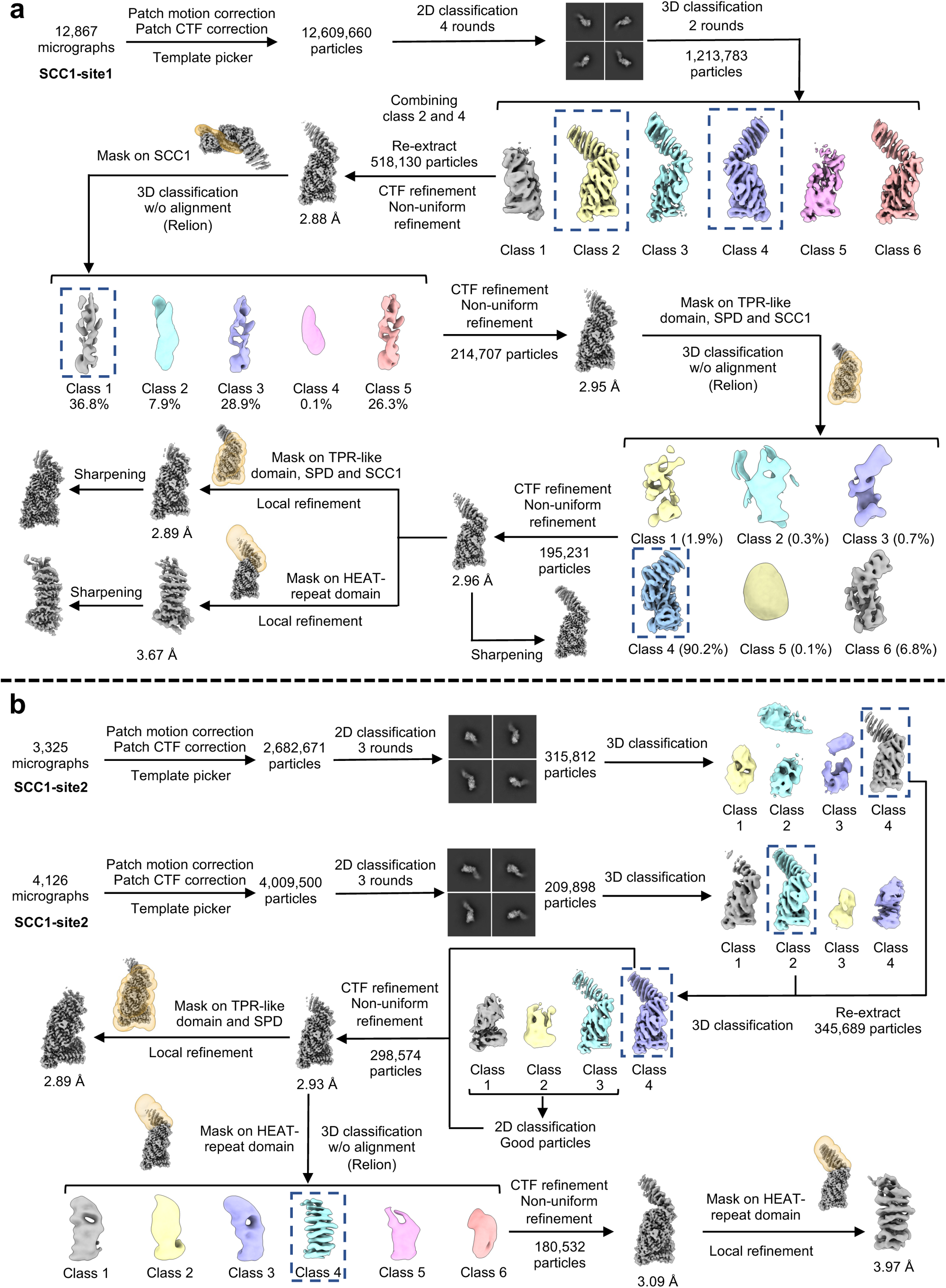
Data-processing flowcharts for separase bound to SCC1^100–320^ or SCC1^310–550^. **a**, CryoEM processing pipeline of SCC1^100–320^–separase^C2029S/ΔAIL1^ complex. After four rounds of 2D classification and two rounds of 3D classification, 518,130 particles were selected and subjected to CTF refinement and non-uniform refinement. 3D classification without alignment was performed to further improve SCC1 density using masks focussed on SCC1 alone and subsequently on SCC1 combined with the C-terminal domains of separase (TPR-like and protease domains). Local refinement with a mask on the HEAT-repeat domain of separase produced a map at 3.67 Å resolution (bottom), improving the density of SCC1 binding to the N-terminal HEAT-repeat domain. **b,** CryoEM processing pipeline of SCC1^310–550^–separase^C2029S/ΔAIL1^ complex. Two data sets (3,325 and 4,126 micrographs) were combined. Following 3D classification 298,574 particles were selected and subjected to CTF refinement, non-uniform refinement, and local refinement. The EM density of HEAT-repeat domain of separase was improved by 3D classification without alignment and local refinement using a mask covering this domain.

**Extended Data figure 8.**
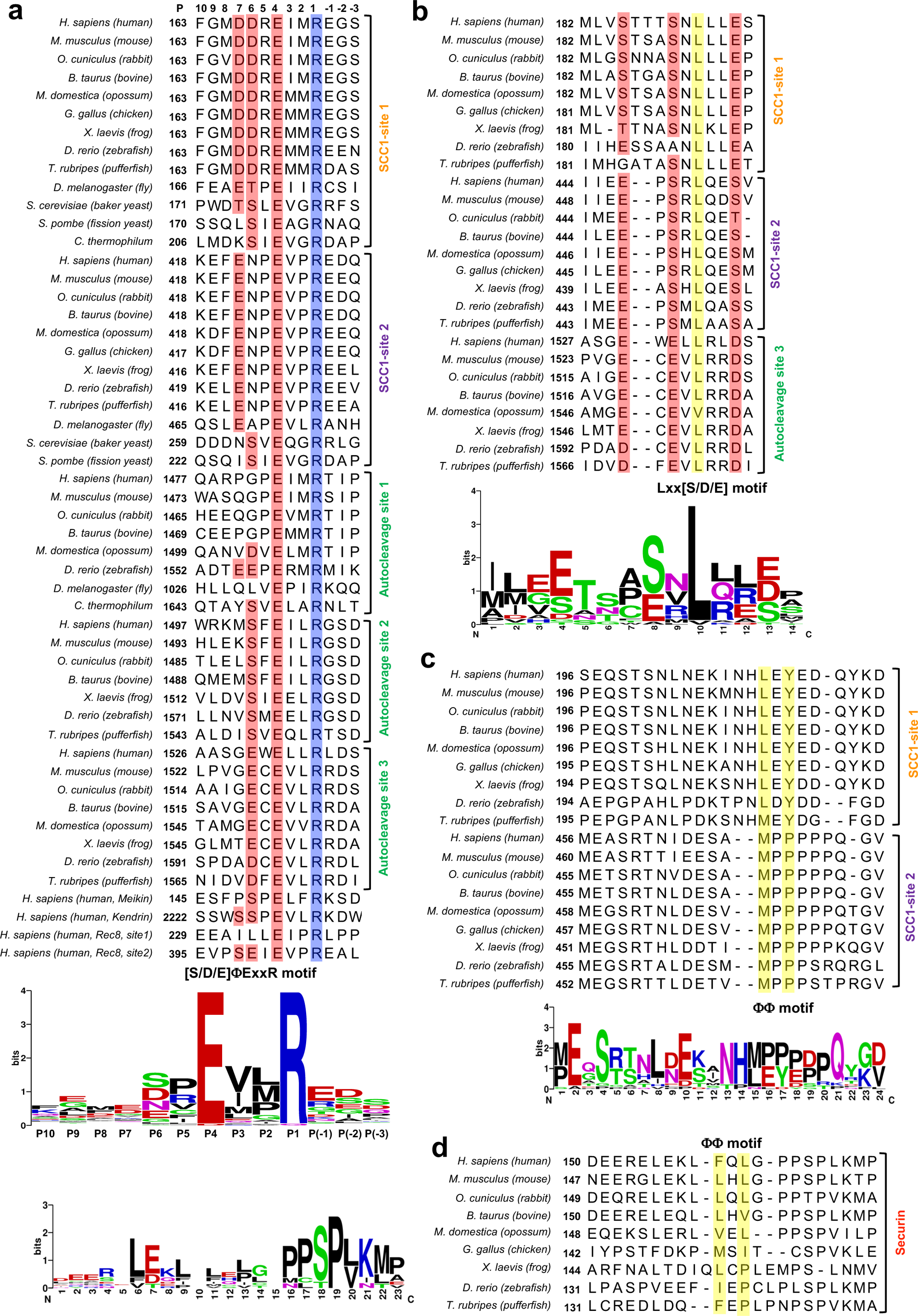
Sequence alignment of substrate motifs. **a**, Sequence alignment of [S/D/E]φExxR motif in SCC1 and autocleavage sites of separase from various species. **b,** Sequence alignment of Lxx[S/D/E] motif in SCC1 and autocleavage site 3 of separase. **c-d,** Sequence alignment of φφ motif in SCC1 and securin. Sequence logos were generated using WebLogo^54^.

**Extended Data figure 9.**
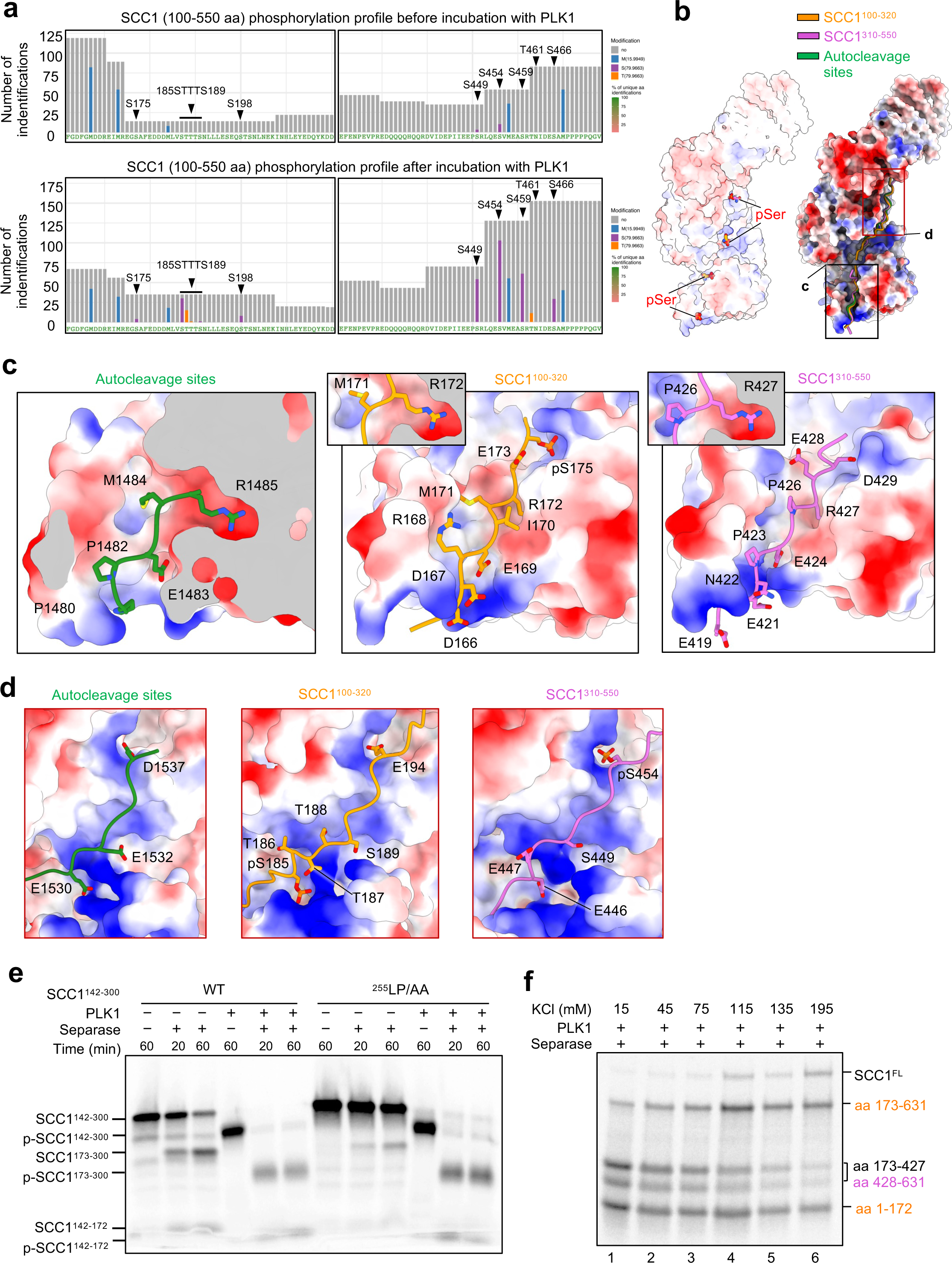
SCC1 binding to separase is primarily mediated by electrostatic interactions. **a**, Mass spectrometry analysis of SCC1 (aa 100-550) after incubation (bottom) with 5 mM ATP, 10 mM Mg^2+^ and 18 ug PLK1 showing the (partially) phosphorylation of residues S175, S185, T186, T187, S189, S449 and S454. **b,** Electrostatic surface potential of the interaction interface between substrates and separase. Left, four phosphate-binding sites on separase. Right, structures of separase^C2029S^ and SCC1^310–550^–separase^C2029S/ΔAIL1^ complex are aligned to the SCC1^100–320^– separase^C2029S/ΔAIL1^ complex. Separase is shown as electrostatic surface representation, while autocleavage sites and SCC1 are shown as cartoons. AIL1 is omitted for clarity to highlight substrate-separase interface. **c,** Close-up view (black box in **b**) showing the autocleavage site 1 and cleavage site motifs binding to the protease domain. The autocleavage site 1 and SCC1 are shown as stick representation. **d,** Close-up view (red box in **b**) showing the autocleavage site 3 and SCC1 exosites binding to the TPR-like domain. **e**, Cleavage assay of SCC1 mutant with alanine mutations in the LPE motif. **f,** Cleavage assay of SCC1 under varying salt concentrations. KCl concentrations ranged from 15 mM to 195 mM, demonstrating the effect of ionic strength on SCC1 cleavage efficiency.

**Extended Data figure 10.**
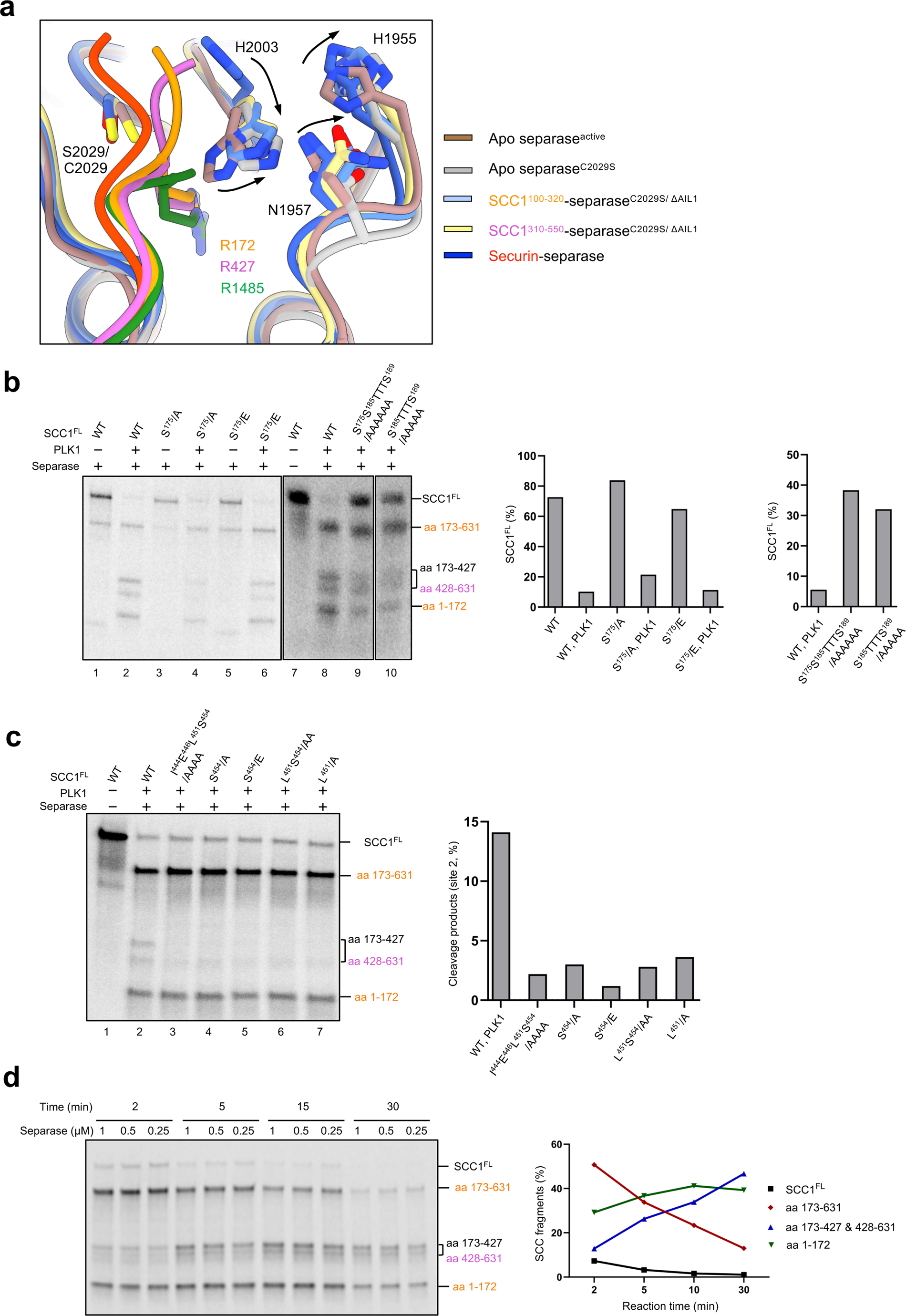
Cleavage assay of SCC1 mutants disrupting the interaction with separase. **a**, Residue arrangement around the catalytic site. The protease domains of apo-separase (active and inactive), securin-separase and SCC1-separase complexes are superimposed and residues around the catalytic site are shown as sticks. Black arrows indicate the movement of H2003, H1955 and N2958 upon substrate binding. **b,** Cleavage assay of SCC1 mutants (phosphorylation sites) near cleavage site 1. Left, autoradiograph of SCC1 cleavage. Right, quantification of uncleaved SCC1. **c,** Cleavage assay of SCC1 mutants near cleavage site 2. Left, autoradiograph of SCC1 cleavage. Right, quantification analysis of the gel. **c,** Cleavage assay of SCC1 mutants (phosphorylation sites) near cleavage site 2. Left, autoradiograph of SCC1 cleavage. Right, quantification of low molecular cleavage products of SCC1. **d,** Cleavage assay of wild-type SCC1 at different concentrations and time points. Left, autoradiograph of SCC1 cleavage. Right, quantification analysis of low molecular weight fragments from different time points at a concentration of 0.5 µM separase.

**Extended Data figure 11.**
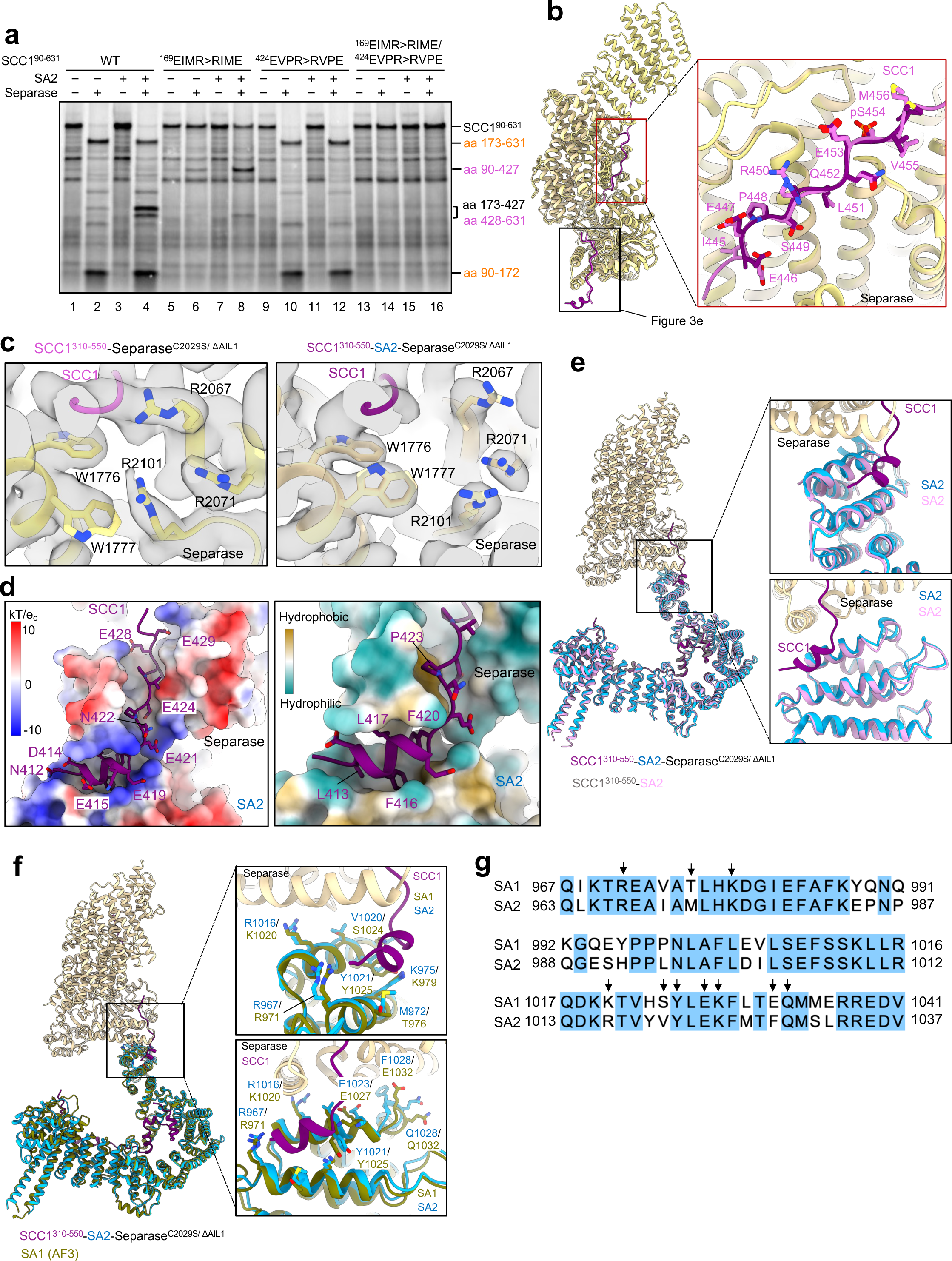
SA2 stimulates the cleavage of SCC1 at site 2. **a**, Cleavage assay of ^35^S-labelled wild-type SCC1 (90-631 aa) and mutants. SCC1 was not phosphorylated but SA2 protein was added in the reactions as indicated. SCC1 mutants for cleavage sites 1 and 2 were generated by swapping the negatively charged glutamate and positively charged arginine in the conserved ExxR motif. **b,** Structural comparison between SCC1^310–550^–SA2-separase^C2029S/ΔAIL1^ and SCC1^310–550^– separase^C2029S/ΔAIL1^ complexes. The two structures were aligned using separase as a reference and are shown as ribbon representation. Black box, close-up view showing SCC1 binding to the protease domain, shown in Figure 3e. Red box, close-up view showing SCC1 binding to the TPR-like domain of separase. SCC1 is depicted as sticks. **c,** Rotamer conformation changes of separase residues interacting with SCC1. Left, the rotamer conformations of W1777, R2067 and R2101 in the SCC1^310– 550^–separase^C2029S/ΔAIL1^ complex. Right, the rotamer conformations of W1777, R2067 and R2101 in the SCC1^310–550^–SA2-separase^C2029S/ΔAIL1^ complex. The corresponding EM densities are shown in grey. **d**, Electrostatic surface potential and the hydrophobicity of the interaction interface. Separase and SA2 are shown as electrostatic surface or hydrophobic surface representations, while SCC1 is shown as cartoon. Electrostatic potentials are contoured from –10 (red) to +10 kTe-1 (blue). Hydrophobic surface is coloured from cyan (hydrophilic) to brown (hydrophobic). **e,** SA2 retains similar conformations before and after binding to separase. The SCC^310–550^–SA2 complex was aligned to the tertiary complex using SA2 as reference. Black boxes, close-up views of the interaction interfaces. **f,** Residues of SA2 interacting with SCC1 and separase are conserved in SA1. The structure of SA1 (olive) predicted by Alphafold 3 was aligned to the tertiary complex using SA2 as reference. Flexible regions of SA1 were omitted for clarity. Black boxes, close-up views of interaction interfaces between SA2, SCC1 and separase with critical residues shown as sticks. **g,** Sequence alignment of SA1 and SA2 at C-terminal regions. Conservation between the sequences is highlighted in blue, with residues mediating interactions with separase indicated by black arrows.

**Extended Data figure 12.**
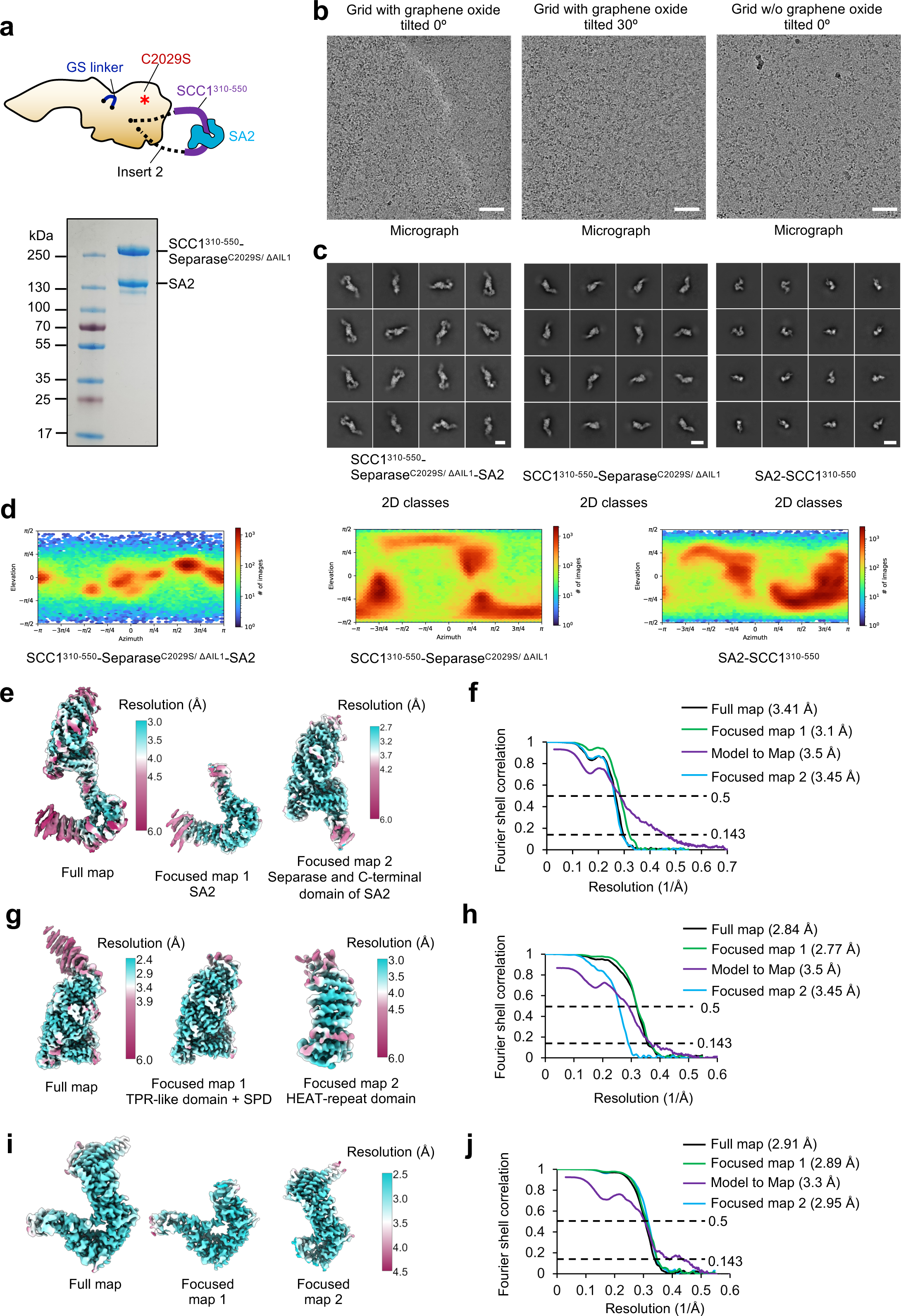
Biochemical and cryoEM analysis of SCC1^310–550^–SA2 complex bound to separase. **a**, Schematic representation and SDS-PAGE gel of the SCC1^310–550^–SA2-separase^C2029S/ΔAIL1^ complex. **b,** Representative cryo-electron micrographs collected on graphene oxide-coated EM grids with the stage untilted and tilted at 30°, and uncoated EM grids without stage tilting. Scale bars, 500 Å. **c,** Gallery of two-dimensional class averages of the tertiary complex (left), SCC1^310–550^– separase^C2029S/ΔAIL1^ complex (middle) and SCC1^310–550^–SA2 complex (right). Scale bars, 100 Å. **d,** Angular distribution plots for the tertiary complex (left), SCC1^310–550^–separase^C2029S/ΔAIL1^ complex (middle) and SCC1^310–550^–SA2 complex (right) calculated using non-uniform refinement algorithm in CryoSPARC^42^. **e-j,** EM density maps of the three complexes colour-coded according to local resolution (**e, g, i**) and gold standard FSC curves for the full map and the focussed refined maps (**f, h, j**). The FSC curves between the full cryoEM map and the final atomic coordinates were calculated using Mtriage^53^.

**Extended Data figure 13.**
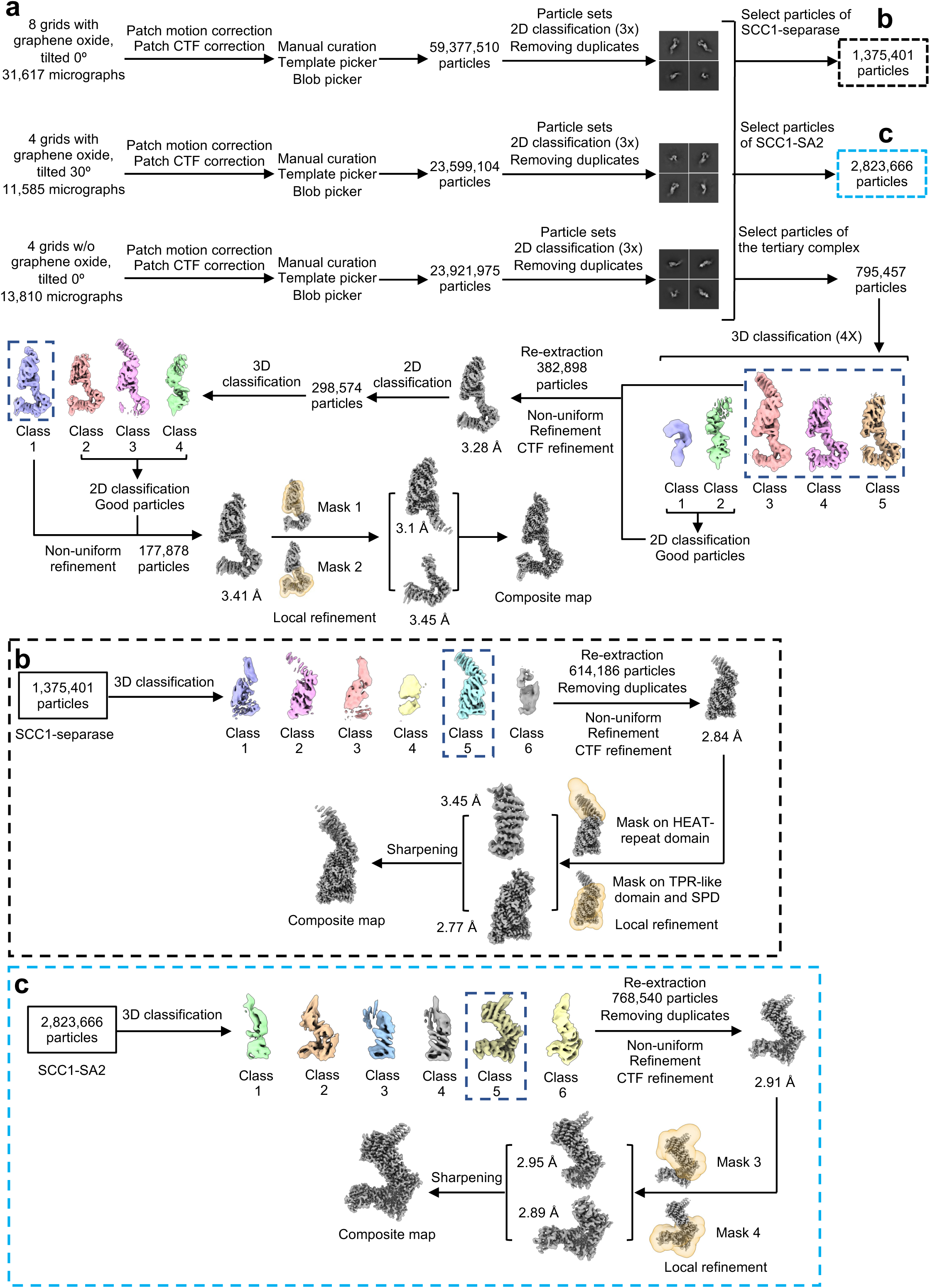
Data-processing flowcharts for SCC1^310–550^–SA2 complex bound to separase. **a**, CryoEM processing pipeline of SCC1^310–550^–SA2-separase^C2029S/ΔAIL1^ complex. Untilted and tilted data sets were initially processed separately. After 2D classification, particles beloniging to the tertiary complex, SCC1-separase and SCC1-SA2 complexes were separated from each other. Particles from each class were selected and combined from all data sets for further processing. Processing of SCC1-separase and SCC1-SA2 complex particles is detailed in panels **b** (black dashed box) and **c** (cyan dashed box), respectively. A total of 795,457 particles of the tertiary complex were subjected to five rounds of 3D classification followed by non-uniform refinement and CTF refinement, producing a map at 3.4 Å resolution. Local refinement using a mask covering SCC1, separase and C-terminal helices of SA2 further improved the density at the interaction interface. A composite map was created by combining two focussed refined maps. **b,** CryoEM processing pipeline of SCC1^310–550^– separase^C2029S/ΔAIL1^ complex resulting in a map of 2.8 Å resolution. Masked refinements on the HEAT-repeat domain and the C-terminal TPR-like and portease domain was used to further improve the EM density. **c,** CryoEM processing pipeline of SCC1^310–550^–SA2 complex. Non-uniform refimement yielded a map a 2.9 Å resolution. Focussed masks are shown on top of the 3D volumes.

**Extended Data figure 14.**
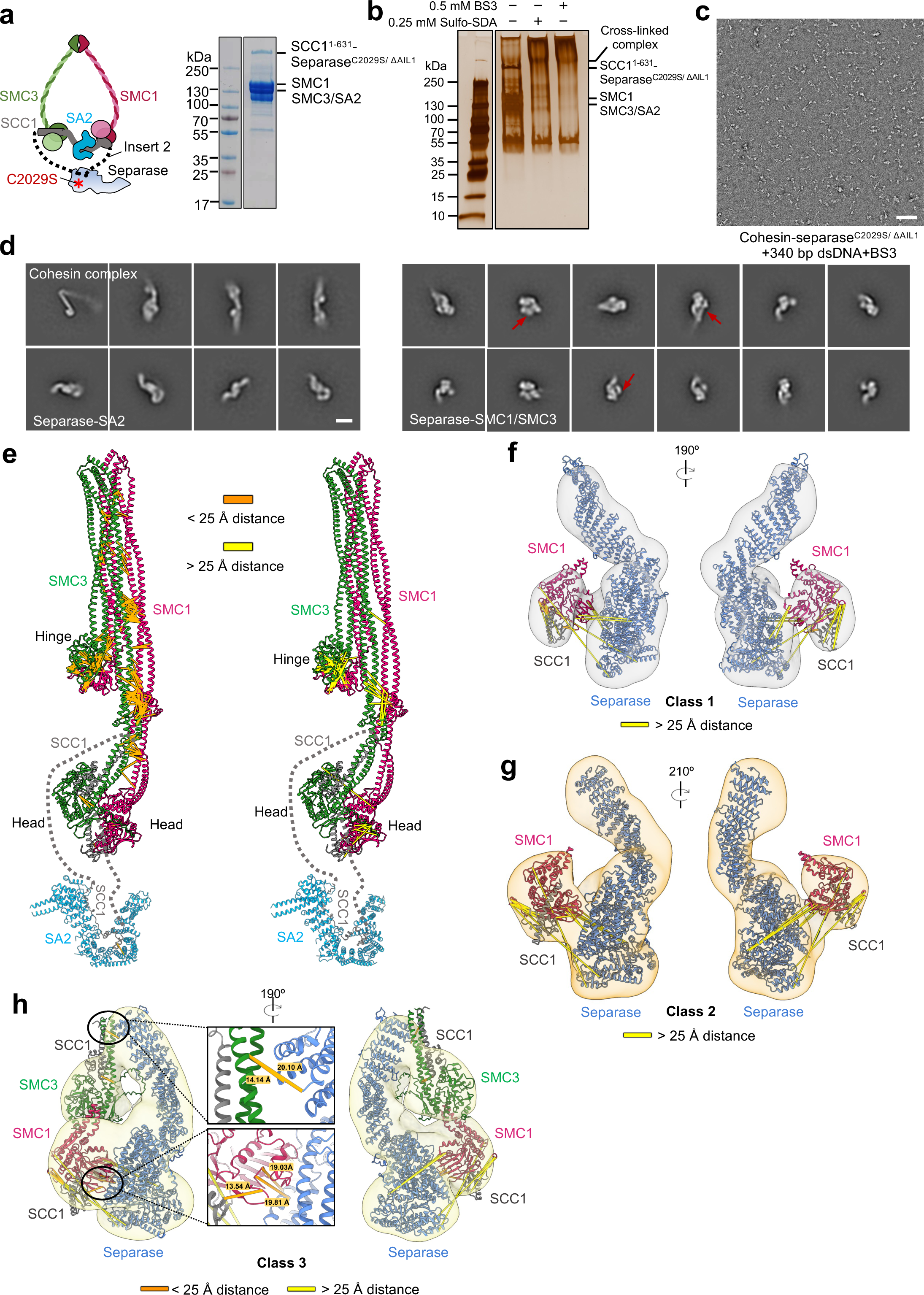
Biochemical and XL-MS analysis of the cohesin-separase complex. **a**, Schematic representation and SDS-PAGE gel of the cohesin-separase complex. **b,** Silver stained SDS-PAGE gel of cohesin-separase complex cross linked with 0.25 mM Sulfo-SDA and 0.5 mM BS3. **c,** Representative negative staining micrograph of cohesin-separase complex incubated with 340 bp double-stranded DNA and 1 mM BS3. Scale bars, 100 nm. **d,** Two-dimensional class averages of the cohesin complex and separase bound to SA2 and potentially SMC subunits. Scale bars, 100 Å. Red arrows indicate the additional densities binding to separase. **e**, Mapping of Sulfo-SDA cross-links with Cα-Cα distances ≤ 25 Å (orange lines) and > 25 Å (gold lines) on the predicted structure of the cohesin complex. The flexible region of SCC1 was omitted and indicated by dashed lines. **f-h**, Three classes of EM maps resulting from negative staining analysis and cross-links mapped on corresponding models. AF3-predicted models of full-length human separase and the head domains of SMC1/3 in complex with SCC1 were fitted into the density maps and shown as cartoon. Cross-links with Cα-Cα distances ≤ 25 Å are shown with orange lines while those > 25 Å are shown with gold lines.

**Extended Data figure 15.**
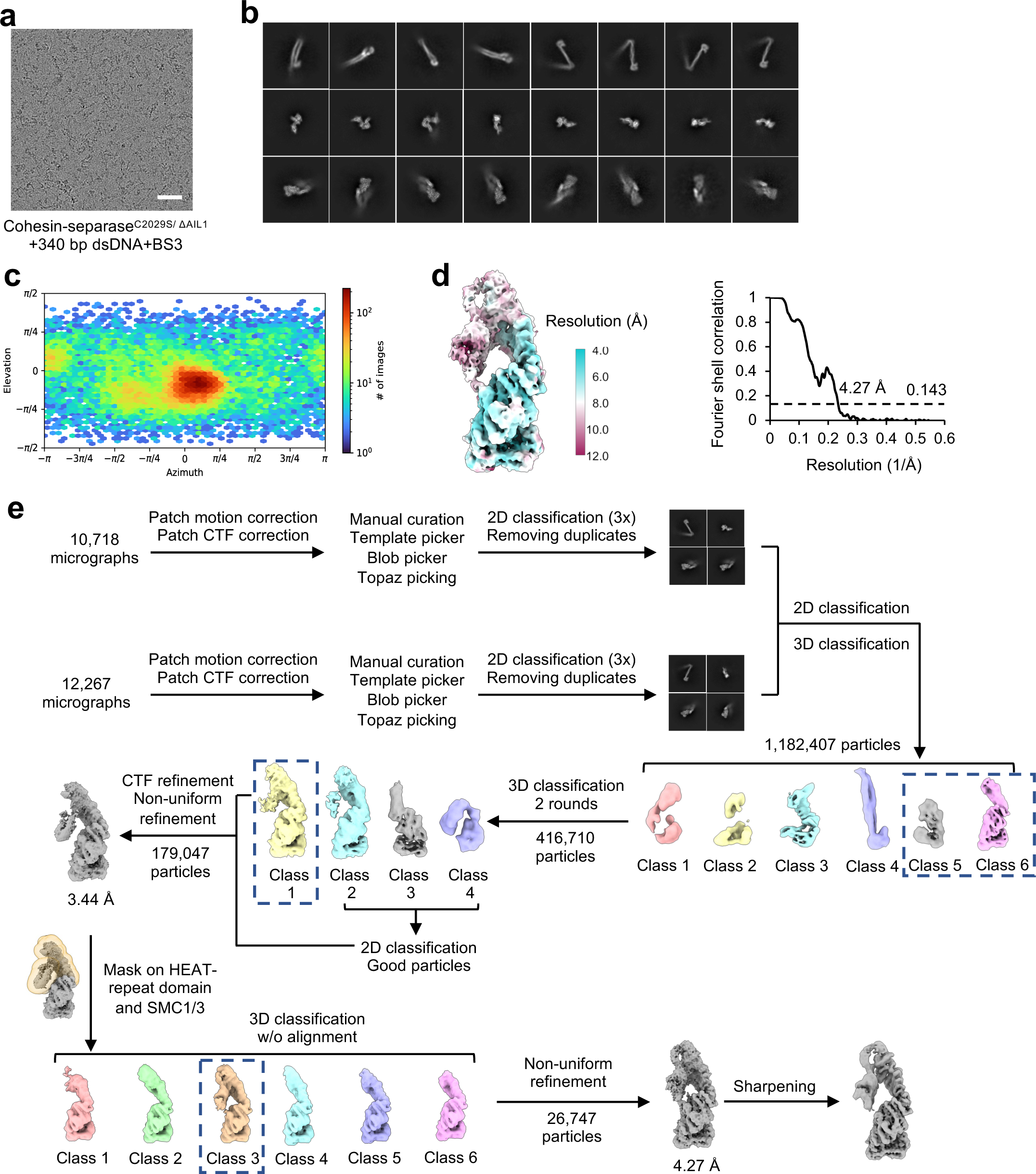
CryoEM analysis of the cohesin-separase complex. **a**, Representative EM micrograph of the cohesin-separase complex incubated with 340 bp double-stranded DNA and cross-linked 1mM BS3. Scale bars, 500 Å. **b,** Representative two-dimensional class averages of the cohesin complex and separase bound to SMC3. Scale bars, 100 Å. **c,** Angular distribution plots for cohesin-separase complex data set calculated using non-uniform refinement algorithm in CryoSPARC^42^. **d,** EM density maps of cohesin-separase complex colour-coded according to local resolution (right) and gold standard FSC curves for the full map (left). **e,** CryoEM processing pipeline of cohesin-separase complex. To improve the density of SMC3 bound to separase, 3D classification without alignment was performed with a mask covering SMC3 and the HEAT-repeat domain of separase.

